# Functional screening of TCR-like antibodies for targeted cancer immunotherapy

**DOI:** 10.1101/2025.07.21.665803

**Authors:** Yi Li, Daosheng Huang, Chang Liu, Zhixiao Zhou, Yilu Song, Dongmei Yang, Wei Rui, Zaopeng Yang, Li Yu, Chenliang Wang, Zheyu Zheng, Jiasheng Wang, Yangxin Fu, Xin Lin

## Abstract

Although hybridomas and display technologies have long been used in antibody discovery, identifying TCR-like antibodies that recognize major histocompatibility (MHC)-presented neoantigens remains an inefficient process. Here, we present a functional screening approach for antibody discovery based on T cell-specific functionality. By constructing <u>S</u>ynthetic <u>T</u> cell receptor and <u>A</u>ntigen <u>R</u>eceptor (STAR)-T cells expressing antibody libraries and co-culturing them with target cells, we observed distinct antigen-specific STAR endocytosis and T cell activation. These two indicators are combined into an E-A functional index, which we demonstrate as an effective screening strategy for identifying antibodies with strict antigen sensitivity and specificity. Using this E-A functional screening method, we successfully discovered antibodies or nanobodies targeting cell-surface tumor antigens and neoantigens presented by human leukocyte antigen (HLA). These antibodies exhibited potent anti-tumor efficacy both *in vitro* and *in vivo*. Moreover, converting these antibodies into chimeric antigen receptors (CARs) and bispecific antibodies also induced T cell activation and tumor killing. In summary, our study establishes a robust functional screening strategy for identifying antibodies or TCR-like nanobodies targeting tumor-specific neoantigens, offering significant potential for advancing targeted cancer therapies.

---

Antibody-based therapies have been developed and improved through decades of biomedical advancement^1^. More than half of the antibody therapeutics are targeting for cancers^2, 3^. Antibodies can target tumor cells by binding to various tumor antigens and exert their therapeutic effects through a variety of mechanisms against tumors^4^. In addition to monoclonal antibodies (mAbs), antibody-drug conjugates (ADCs) and bispecific antibodies (BsAbs), T cell therapies utilizing antibody fragments as targeting moieties have emerged as promising strategies for targeted cancer therapy in recent years, which include CAR-T cell and STAR-T cell therapies^5–7^. In this context, the discovery of antibodies with high affinity and specificity is essential for the development of effective antibody-based T cell therapies for cancer therapy.

Traditional methods for screening specific antibodies primarily involve hybridoma technology and antibody engineering techniques, including phage display antibody libraries, ribosome antibody libraries, yeast surface displays, transgenic mice, and others^8–12^. While these established traditional methods have proven successful in obtaining high-affinity antibodies, shortcomings and limitations exist such as non-specific binding, loss of valuable clones, complex procedures, time intensiveness and high costs. In recent years, novel strategies have emerged to address these drawbacks. Single cell antibody screening methods, such as microfluidic technology-based antibody library screening, have gained attention for their ability to identify rare and desirable antibody clones due to their high-throughput and rapidity^13–17^. However, these systems necessitate specialized equipment and expertise, contributing to their high cost. With the advancement of artificial intelligence, structure-based and sequence-based deep leaning networks have been explored for training in the design of novel antibodies, such as the variable domain of heavy chain of heavy-chain-only antibody (HCAb), also known as VHH or nanobody. However, the success rates of these attempts are relatively low and the binding affinities of the designed antibodies are modest^18^. Instead of relying solely on screening by binding affinity of antigen-antibody complex, functional screening utilizing antibody-specific properties may offer a more efficient approach. This method could potentially reduce the need for additional rounds of functional validation and directly select functional antibodies for T cell therapies. An innovative approach involves utilizing CAR-T cells to screen single-chain fragment variable (scFv) antibodies, leveraging T cell activation properties as a cell-based functional selection strategy^19, 20^. Although these approaches provide valuable insights, they are challenged by limitations such as the high tonic signals of CAR-T cells. Thus, there remains a need for more suitable screening indicators and more effective functional screening strategies for antibody discovery.

T cell receptor (TCR) endocytosis has been reported to play various roles in regulating T cell functions such as TCR signaling, which is considered crucial for modulating T cell activation^21, 22^. CD69 is a well-known classical early marker of lymphocyte activation^23^. In the present study, we observed robust endocytosis of STAR receptor carrying antibody fragments scFv or VHH, and CD69 activation in an antigen-specific manner in STAR-T cells. Based on these, we designed a novel functional screening approach by using both STAR molecule endocytosis and CD69 activation as functional readouts. This approach enables rapid isolation of antigen-specific antibody fragments from antibody-based STAR-T cell libraries. Using this method, we successfully identified several VHHs targeting tumor cell surface antigen such as CD22 and, importantly, several VHHs targeting tumor frequent hotspot neoantigen P53^R175H^ presented by HLA-A*02:01 promisingly. The STAR-T cells engineered with identified VHHs exhibited potent anti-tumor efficacy both *in vitr*o and *in vivo*. Furthermore, the same functionality extends to other therapeutic modalities, such as CAR-T and BsAb. These findings introduce a novel strategy for identifying TCR-like antibodies targeting a wide range of novel neoantigens, offering more efficient steps, shorter timelines, and higher output ratios for cancer immunotherapy.

## Results

### Membrane antigen triggers antigen-specific endocytosis and activation of STAR-T cells

As a novel double-chain TCR-based chimeric receptor, STAR combines antigen-recognition domain of antibody with constant regions of TCR that engaging endogenous CD3 signaling machinery, which has been reported to trigger low tonic signaling than CAR (**Supplementary Fig. 1a**)^5–7^. STAR-T cells exhibit higher antigen sensitivity and superior anti-tumor effects than CAR-T cells, making them the preferred carrier in our study. We first established cell lines expressing cognate antigen-STAR pairs, which included K562 cells expressing the membrane antigen mesothelin (MSLN) or CD123 and a TCR-deficient Jurkat cell line, JC5, expressing their cognate STARs with MSLN VHH or CD123 VHH. To search for applicable indicators of T cell functions upon antigen-specific stimulation, we co-incubated K562 cells with their cognate or non-cognate VHH STAR-JC5 cells and assessed expression levels of surface molecules on JC5 cells using flow cytometry by immunostaining (**Fig. 1a, Supplementary Fig. 1b** and 2a). Notably, we observed complete conversion of the membrane receptor STAR on JC5 cells from positive to negative by staining of TCRα/β antibody, and the classical lymphocyte activation marker CD69 was upregulated in JC5 cells following stimulation with their cognate antigens after 24 hours of co-culture (**Fig. 1b and Supplementary Fig. 2b**). To confirm the universality of these findings, we repeated the experiment using cognate antigen-scFv STAR pairs, involving K562 cells expressing the membrane antigen CD19 or glypican 3 (GPC3), and JC5 cells expressing their cognate scFv STARs, which showed similar results to the VHH groups (**Fig. 1c, d and Supplementary Fig. 2c**, d). Thus, our results indicate that STAR-JC5 cells exhibit antigen-specific STAR endocytosis and CD69 activation.

**Fig. 1.**
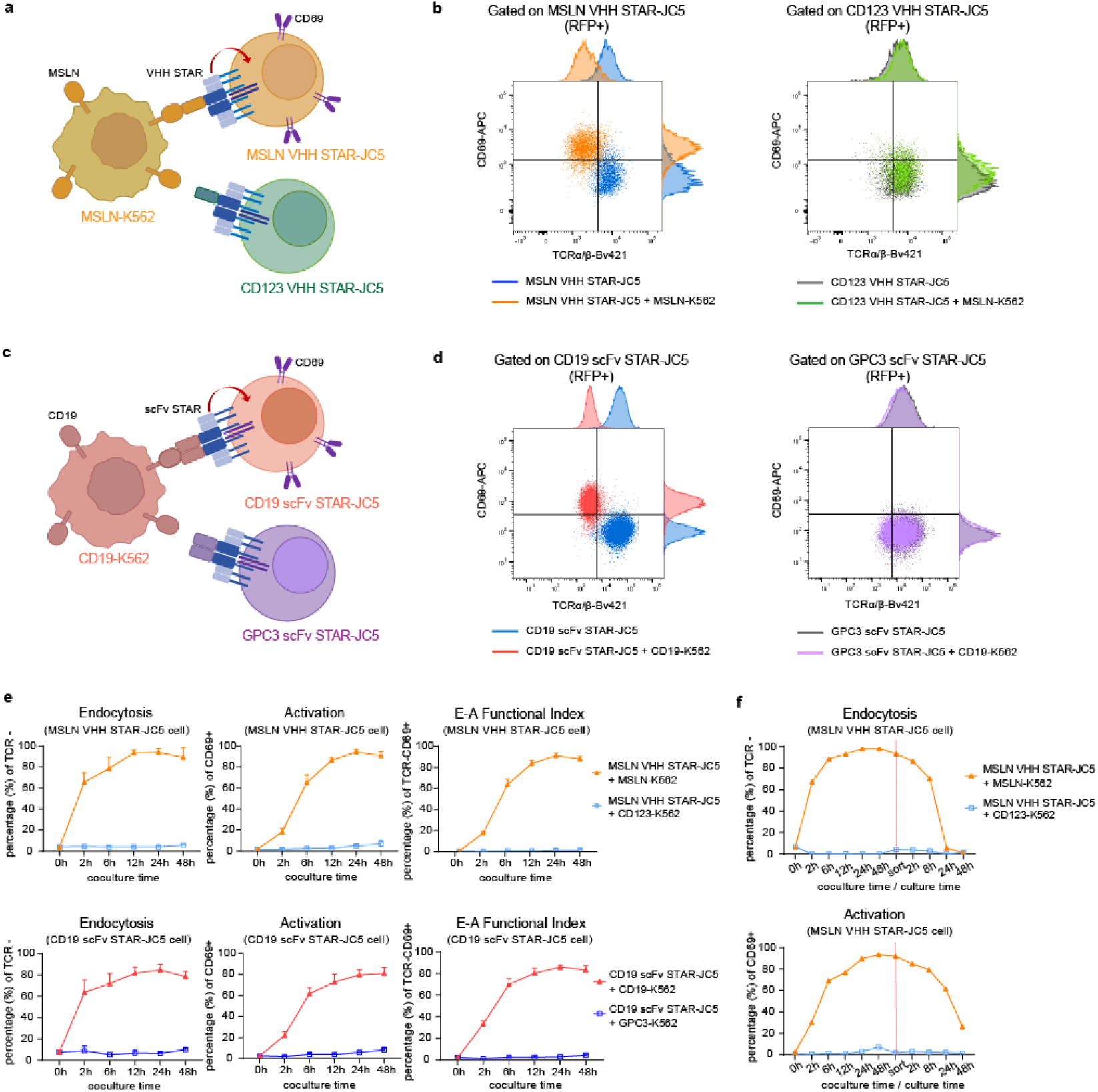
STAR endocytosis and CD69 activation of T cells are membrane antigen-specific. **a, c**, Schematic of the functional evaluation assay of the VHH STAR-JC5 cells and scFv STAR-JC5 cells. For VHH STAR group (a), K562 target cells were transduced to express tumor membrane antigen MSLN and subsequently co-cultured with JC5 cells transduced with cognate MSLN VHH STAR or non-cognate CD123 VHH STAR. For scFv STAR group (c), K562 target cells were transduced to express tumor membrane antigen CD19 and subsequently co-cultured with JC5 cells transduced with cognate CD19 scFv STAR or non-cognate GPC3 scFv STAR. Same coloring indicates cognate antigen-antibody pairs. Red arrows represent endocytosis process. Purple homodimers represent surface molecular CD69. **b, d**, The expressions of STAR and CD69 in RFP positive STAR-JC5 cells were detected by staining with antibodies specific to human TCRα/β and CD69 after co-culture for 24h. **e**, Flow cytometry analysis of STAR endocytosis (TCRα/β negative), CD69 activation (CD69 positive) and E-A functional index (TCRα/β negative and CD69 positive) in RFP positive STAR-JC5 cells after co-incubation with target K562 cells expressing cognate or non-cognate membrane antigen at different time points. **f**, Flow cytometry analysis of STAR-JC5 cells endocytosis and activation at different time point before and after JC5 cells were sorted at 48h of co-culture by FACS. The E:T ratio used in co-culture experiments is 1:1. Data in (**b**) and (**d-f**) are representative of three independent experiments. Data in (**e**) are presented as mean ± SEM.

We further detected STAR endocytosis and CD69 activation of JC5 cells at different time points after co-culture with target cells. We found that STAR endocytosis occurred more rapidly than CD69 expression within the first two hours and these two indicators exhibited strict synchronicity over time (**Fig. 1e and Supplementary Fig. 2e**). Based on above findings, we integrated endocytosis and activation as a functional readout termed “E-A functional index”, which represents TCR-negative and CD69-positive (TCR^-^CD69^+^) states. This dual-readouts strategy effectively mitigates the enrichment of spontaneously activated clones that may arise from using CD69 as single-readout, while simultaneously eliminating interference from STAR-negative clones that could arise when relying solely on TCR endocytosis, thereby enhancing the efficiency and accuracy of antibody selection (**Supplementary Fig. 3**). Upon removal of specific antigenic stimulation after 48 hours of co-culture, we found the endocytosed STAR receptors rapidly recycled to the T cell surface within 24 hours, while the activated CD69 reverted to a resting state in a short period (**Fig. 1f and Supplementary Fig. 2f**). These findings suggest that the E-A functional index is recyclable and can be re-used to assess T cells effector function. In addition, we expressed mutSTAR in wild-type (wt) Jurkat cells confirming that TCR endocytosis is directly linked to specific antigen binding (**Supplementary Fig. 4**).

### E-A functional screening enables isolation of cognate membrane antigen-specific antibodies

To evaluate sensitivity of E-A functional index, we firstly co-incubated MSLN VHH STAR-JC5 cells with cognate MSLN-K562 cells at various effector-to-target (E:T) ratios ranging from 1000:1 to 1:10, finding that the functional indicators scaled proportionally with the density of specific antigens, with only 2% of cognate target cells capable of triggering over 50% endocytosis and activation of JC5 cells (**Supplementary Fig. 5a**, b). Furthermore, we determined whether the E-A functional index is sufficiently sensitive for identifying a cognate antibody from a library of non-cognate antibodies. We firstly generated three proof-of-concept libraries of MSLN VHH STAR-JC5 cells with 1:1, 1:100, and 1:10,000 mixtures of other STAR-JC5 cells (**Fig. 2a**). The E-A functional index was detected after co-incubation of these mixtures with wt K562 cells or target MSLN-K562 cells (**Fig. 2b**). As expected, the E-A functional index (TCR^-^CD69^+^ %) of JC5 cells was approximately 50% in the 1:1 group, while the percentages were 2.46% in the 1:100 group and 0.8% in the 1:10,000 group co-cultured with target MSLN-K562 cells.

**Fig. 2.**
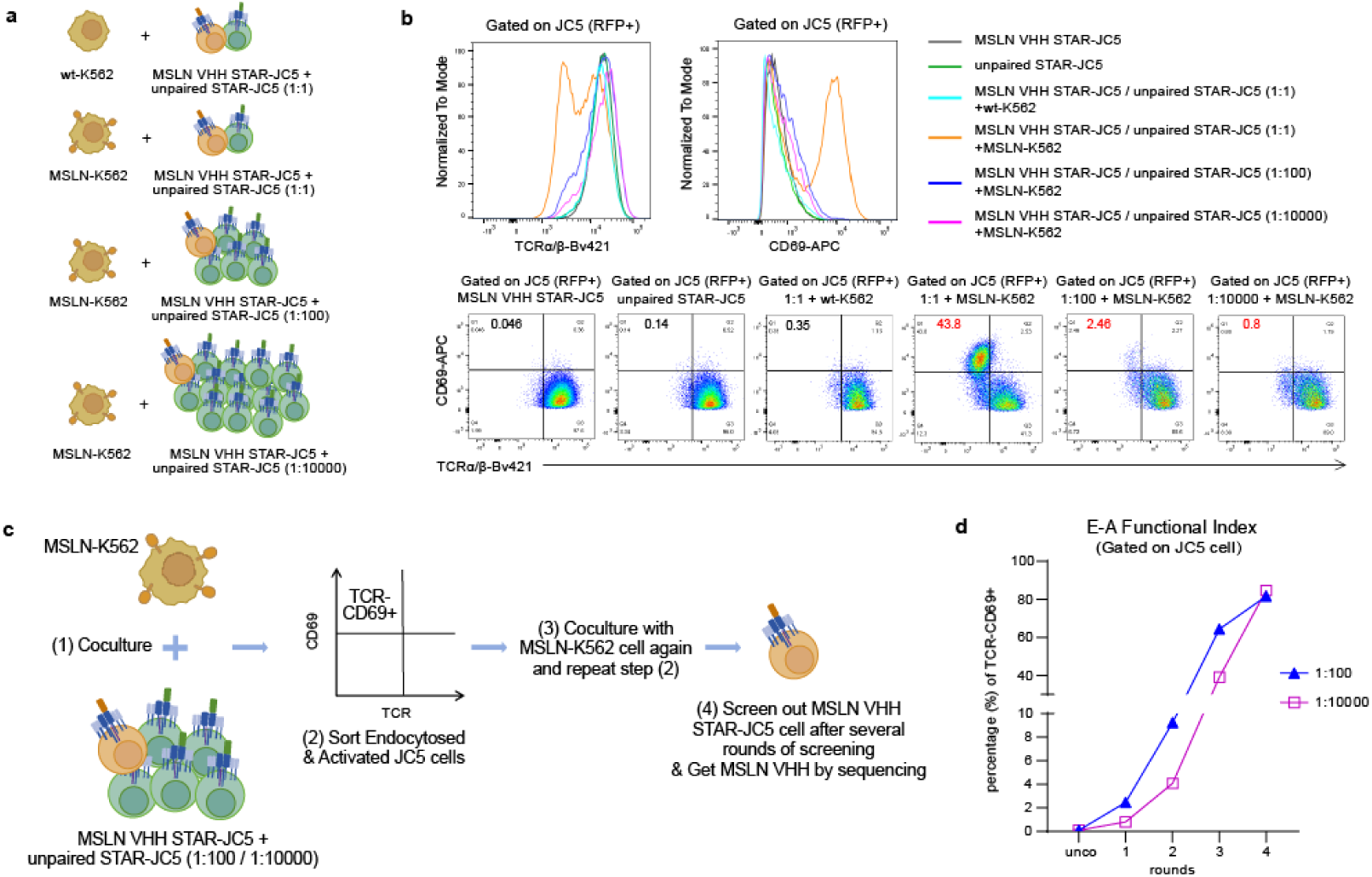
E-A functional index is sufficiently sensitive to identify one cognate antibody from 10,000 non-cognate antibodies library. **a, b**, Schematic of the sensitivity evaluation assay and flow cytometry analysis of endocytosis and activation for the mixture of cognate MSLN VHH STAR-JC5 cells and non-cognate STAR-JC5 cells co-cultured with wt-K562 cells or target MSLN-K562 cells. **c**, Schematic outline of the functional screening method based on E-A functional index to isolate one cognate antibody from 100 or 10,000 non-cognate antibodies libraries. **d**, Flow cytometry analysis of E-A functional index in gated JC5 cells of 1:100 or 1:10,000 mixtures after each round of E-A functional screening. Data in (**b**) are representative of three independent experiments.

Based on these results, we developed an antibody screening strategy based on E-A functional index termed E-A functional screening. We firstly used E-A functional screening to isolate MSLN VHH STAR-JC5 cells from the 1:100 and 1:10,000 libraries by fluorescence-activated cell sorting (FACS) and subsequently identified sequences of enriched antibodies through sequencing (**Fig. 2c**). Following four rounds of E-A functional screening, the TCR^-^CD69^+^ JC5 cells were notably enriched to over 80% in both the 1:100 and 1:10,000 groups and were sorted for Sanger sequencing (**Fig. 2d and Supplementary Fig. 5c**). The sequencing results confirmed the enrichment of the expected MSLN VHH antibody, with almost no other sequence peaks detected (sequencing results not shown). Therefore, the E-A functional index can effectively isolate cognate antibodies carried on STAR-JC5 cells even at a ratio as low as 1:10,000.

To further validate the sensitivity and specificity of E-A functional screening, we constructed a VHH library by immunizing alpacas with CD22 protein and subsequently engineered VHH into STAR structure. The estimated diversity of this VHH STAR library is over 10^6^. The VHH STAR library was then transduced into JC5 cells and co-cultured with K562 cells expressing the target antigen CD22 (**Fig. 3a**). Functional screening against CD22 was conducted over four rounds until the E-A functional index reached 50% in the co-culture group (**Fig. 3b and Supplementary Fig. 6a**). The TCR^-^CD69^+^ JC5 cells obtained after the fourth round of screening were sorted using FACS and subjected to NGS sequencing as the output to identify enriched VHHs, while the original pooled CD22 VHH library served as the input (**Fig. 3c**). The NGS sequencing results indicated that 5 of the top 10 enriched VHHs shared the same CDR3 regions (CDR3 of top 1, 2 and 6: DPSFDTPWYRYAY, CDR3 of top 4 and 8: GVLSVCELTDSYQY). We then picked up the top 10 enriched VHHs and constructed them into STAR-JC5 cells, and their E-A functional index was re-evaluated upon co-incubation with CD22-K562 cells, demonstrating substantial endocytosis and activation for 7 of the top 10 CD22 VHHs STAR-JC5 cells (except CD22-5, 7, 9), especially the five VHH groups with shared CDR3 regions (**Fig. 3d and Supplementary Fig. 6b**, c). To further confirm the functional efficacy of the selected seven antibodies, primary T cells engineered with these seven CD22 VHH mutSTARs were generated (mutSTAR being a mutant of STAR with modifications enhancing surface display efficiency and antigen-binding ability^5^). These cells were then co-cultured with two target tumor cell lines expressing luciferase (luc): human B lymphoblastoid Raji cells which endogenously express CD22, and CD22-knockout Raji cells as a negative control. Consistently, the seven CD22 VHH mutSTAR-T cells exhibited robust CD22-specific killing effects on tumor cells and cytokines (IFN-γ, TNF-α, IL-2) secretion capacity *in vitro* (**Fig. 3e, f**). We further evaluated the binding affinity using surface plasmon resonance (SPR), which revealed that the CD22-1 VHH bound to human CD22 protein with a dissociation constant (Kd) of 3.55 nM (**Supplementary Fig. 6d**).

**Fig. 3.**
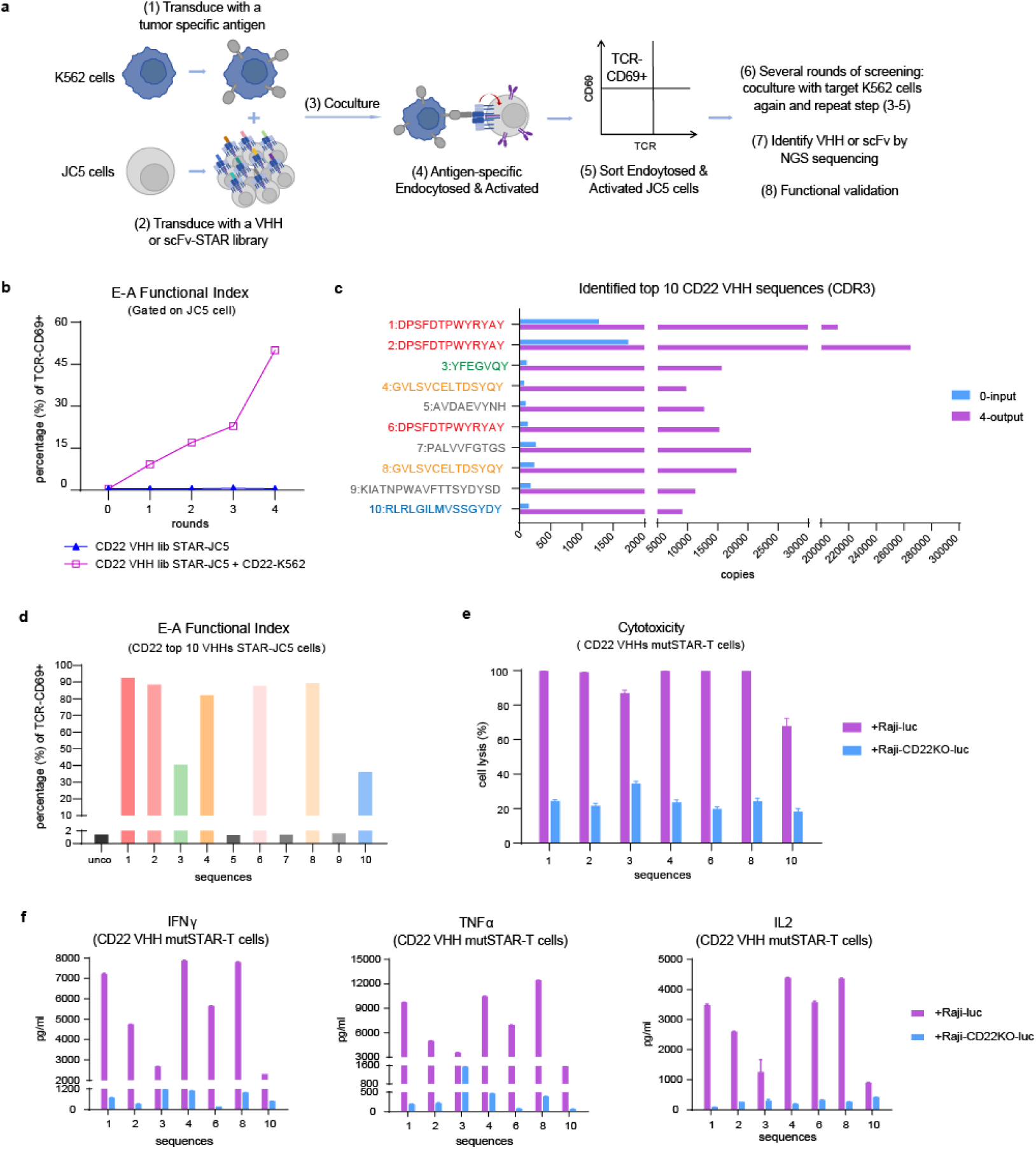
E-A functional screening identifies antibodies targeting CD22. **a**, Schematic diagram of the technical route of the E-A functional screening approach used for tumor specific antigen-specific antibodies screening. **b**, Flow cytometry analysis of E-A functional index in gated CD22 VHH lib STAR-JC5 cells after each round of E-A functional screening. **c**, Identification of enriched CD22 VHHs by NGS (CDR3 regions amino acid sequences of VHHs are shown). 0-input represents original CD22 VHH lib. 4-output represents CD22 VHHs screened after four rounds of functional selection. Top ten enriched CD22 VHHs are ranked by ratio of 4-output over 0-input. **d**, Flow cytometry analysis of E-A functional index in top ten enriched CD22 VHHs-based STAR-JC5 cells co-cultured with target CD22-K562 cells. **e**, Cytotoxicity (presented as percentage of specific cell lysis) of seven E-A functional index-positive CD22 VHHs-based STAR-T cells against Raji cells expressing luciferase (Raji-luc) and CD22 knockout Raji-luc cells (Raji-CD22KO-luc). **f**, The supernatants from (e) co-culture were analyzed by ELISA for secreted IFN-γ, TNF-α and IL-2. The co-culture time used in the experiments is 24h and the E:T ratio is 1:1. Data in (**d-f**) are representative of three independent experiments. Data in (**e, f**) are presented as mean ± SEM.

Next, we selected CD22-1 VHH-based mutSTAR-T cells to determine whether it could control tumor growth *in vivo*. We first engrafted Raji-luc cells into NOD-Prkdc^em26Cd52^Il2rg^em26Cd22^/NjuCrl (NCG) mice through intravenous injection (**Fig. 4a**). The mice were divided into two groups according to luminescence quantification of tumor burden, and the CD22-1 VHH mutSTAR-T cells were infused intravenously at day 4. Mock-T cells with exogenous RFP transduction were infused as negative controls. The tumor burden were monitored every few days. We found that the tumor growth was markedly suppressed and the survival rate of the mice was significantly improved in the CD22-1 VHH group (P<0.0001) (**Fig. 4b-d**). The *in vivo* experiments indicate that the novel P53 VHH-based STAR-T cells exhibit potent anti-tumor efficacy. By contrast, we established Raji-CD22KO-luc tumor engrafted mouse model as a control to evaluate the off-target effects and *in vivo* toxicity of the CD22-1 VHH, founding that the CD22-1 VHH mutSTAR-T cells had no effect on Raji-CD22KO-luc tumors (**Fig. 4e-h**). These findings demonstrate the effectiveness of the E-A functional screening as a robust approach to identify cognate antibodies targeting tumor membrane antigens for cancer immunotherapy.

**Fig. 4.**
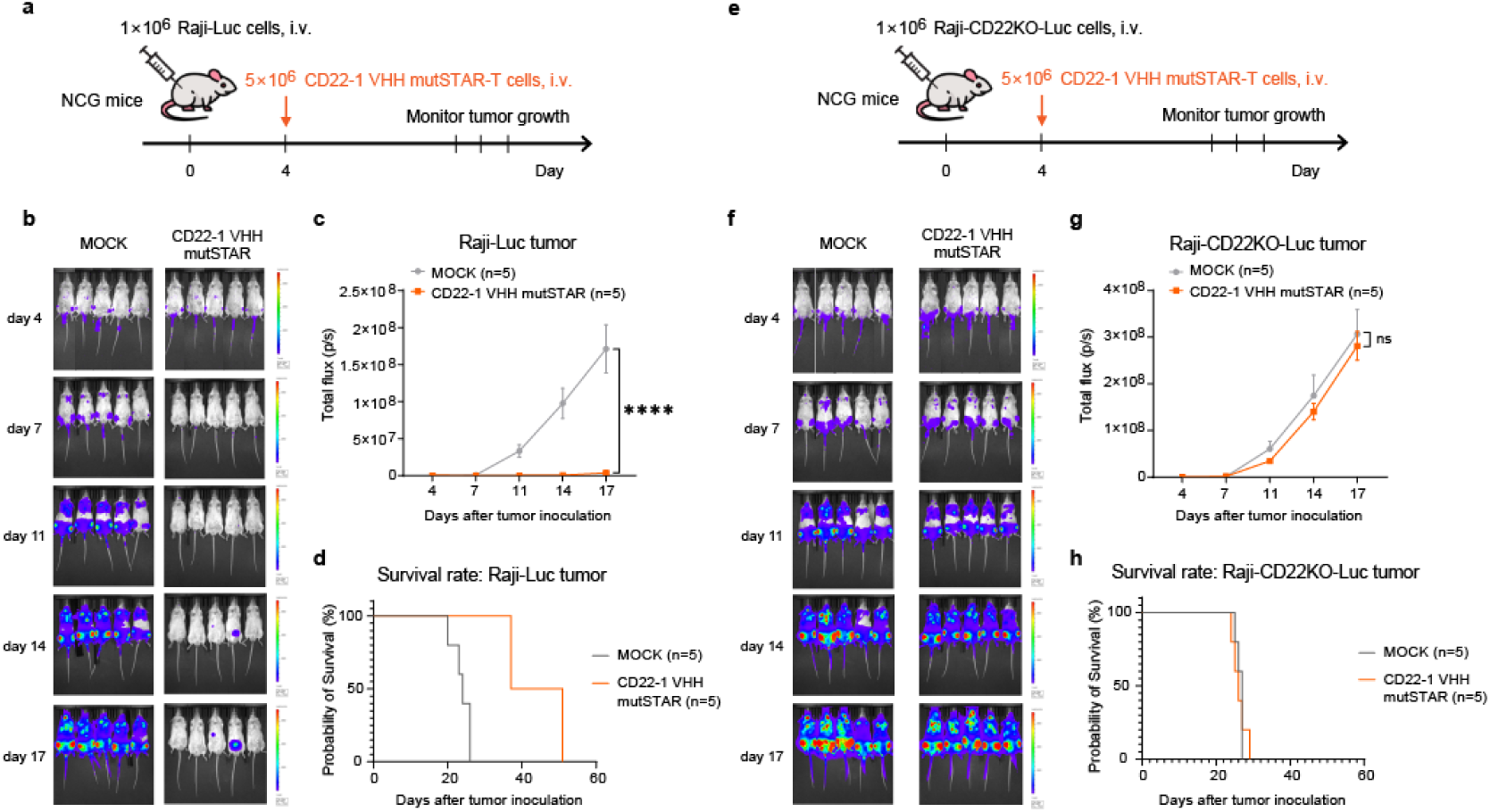
In vivo anti-tumor efficacy of CD22-1 VHH mutSTAR-T cells. **a, e**, Schematic of the experimental process for *in vivo* mouse model xenografted with Raji (lymphoma) tumor cells. NCG mice were inoculated intravenously with 1×10^6^ Raji-Luc cells endogenously express CD22 (**a-c**) or Raji-CD22KO-Luc cells (**e-g**) at day 0, and 5×10^6^ CD22-1 VHH mutSTAR-T cells were intravenously injected on day 4. Tumor growth was monitored by *in vivo* luminescence imaging. **b, f**, Bioluminescence images of tumors. MOCK group were infused with T cells expressing RFP. **c, g**, Tumor progression as evaluated by bioluminescence imaging. **d, h**, Survival rate curves of the xenografted mice (N=5 per group). Statistical analysis of quantification in (**c, g**) was performed using two-way analysis of variance (ANOVA). Data are representative of three independent experiments and are presented as mean ± SEM at the last time point. P values were presented as indicated. ****P<0.0001, ns means no significant difference (P>0.05).

### E-A functional screening identifies novel neoantigen-specific TCR-like antibodies

The membrane surface of tumor cells harbors numerous antigen molecules that can be targeted by immune cells. These antigens encompass not only tumor-associated membrane antigens, but also tumor-specific neoantigens generated by genomic alteration of tumor cells, such as CDK4^R24C^, KRAS^G12V/C/D^, and others, which can be presented on the membrane surface by HLA molecules^24^. After confirming the feasibility of the E-A functional screening strategy for identifying membrane antigen-specific antibodies, we aimed to determine whether this strategy could be extended for identification of neoantigen-specific antibodies. We selected the most frequent missense mutant of tumor-suppressor gene TP53 (P53^R175H^, presented by HLA-A*02:01) as a target neoantigen, and its specific scFv (H2)^25^ was constructed into STAR-JC5 cells as a positive control for coculture experiments. Meanwhile, we pulsed K562 cells expressing HLA-A*02:01 (K562-A2) and T2 cells that express HLA-A*02:01 with P53 R175H mutant (mut) peptide (<u>HMTEVVR**H**C</u>) or P53 wildtype (wt) peptide (<u>HMTEVVRRC</u>) at various concentrations as target cells. After co-incubation, we found that only mut peptide-pulsed target cells triggered endocytosis and activation of P53 scFv STAR-JC5 cells, exhibiting a peptide-concentration-dependent increase (unlike in the wt peptide-pulsed groups) (**Fig. 5a**). Moreover, T2 cells performed superior stimulation effects then K562 cells, probably due to T2 cells being TAP-deficient, allowing them to present purer exogenous peptides and thus serving as a more suitable carrier. These results suggest that the E-A functional index is neoantigen-specific, and its magnitude depends on antigen density.

**Fig. 5.**
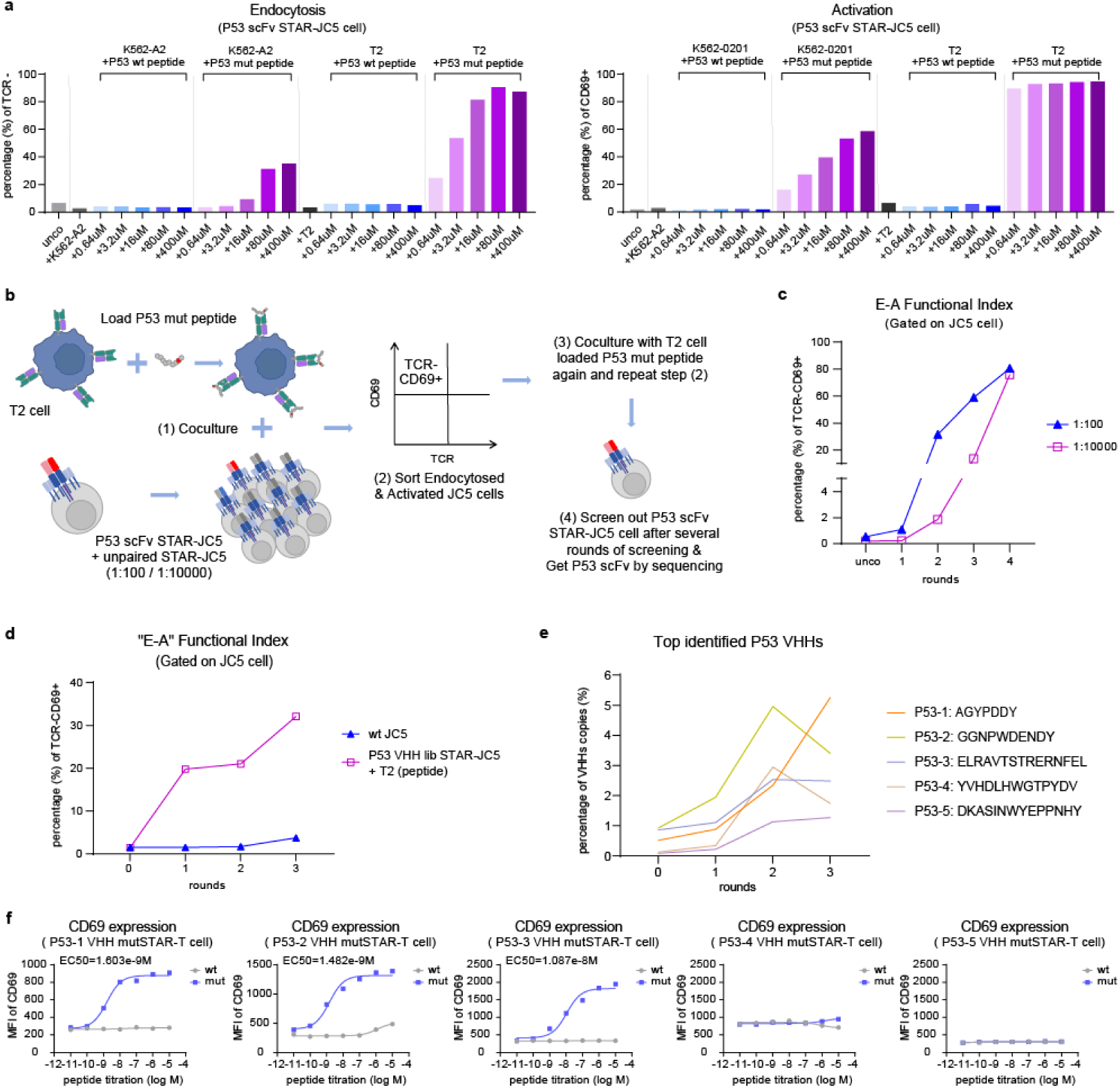
E-A functional screening enables identification of neoantigen P53^R175H^-specific antibodies. **a**, Flow cytometry analysis of STAR endocytosis and CD69 activation in P53 scFv STAR-JC5 cells co-cultured with target K562 cells expressing HLA-A*02:01 (K562-A2) or T2 cells loaded with different doses of P53 wt peptide (HMTEVVRRC) or P53 mut peptide (R175H: HMTEVVRHC). **b**, Schematic outline of the E-A functional screening method to isolate one cognate neoantigen-specific antibody from 100 or 10,000 non-cognate antibodies libraries. **c**, Flow cytometry analysis of E-A functional index in gated JC5 cells of 1:100 or 1:10,000 mixtures after each round of functional screening co-cultured with T2 cells loaded with P53 R175H mut peptides. Peptides concentration used in each round is 10 µM. (d) Flow cytometry analysis of E-A functional index in gated P53 VHH lib STAR-JC5 cells before and after three round of functional screening. Target cells used in the first and third rounds of screening are T2 cells loaded P53 R175H mut peptides and the second round used P53 wt peptides. Peptides concentration used in each round is 1 µM. **e**, Top five enriched P53 VHHs identified by NGS (presented as percentage of VHHs copies in each round of functional screening, CDR3 regions amino acid sequences of VHHs are shown). **f**, Flow cytometry analysis of CD69 activation in top five enriched P53 VHHs-based mutSTAR-T cells co-cultured with T2 cells loaded with different doses of P53 wt peptide or R175H mut peptide. MFI, mean fluorescence intensity. EC50, half maximal effective concentration. Data in (**a, f**) are representative of three independent experiments.

To ascertain the sensitivity of the E-A functional index in isolating neoantigen-specific antibodies, we repeated the 1:100 and 1:10,000 experiments as previously described, utilizing P53 scFv STAR-JC5 cells mixed with other STAR-JC5 cells for co-culture with T2 cells loaded with cognate P53 R175H mut peptides (**Fig 5b**). After four rounds of functional screening, the JC5 cells positive for E-A functional index were substantially enriched to approximately 80% in both the 1:100 and 1:10,000 groups. Sequencing results confirmed the enrichment of the correct P53 scFv antibody (**Fig 5c and Supplementary Fig. 7a**, sequencing results not shown). Thus, the E-A functional index can effectively isolate cognate neoantigen-specific antibodies carried on STAR-JC5 cells even at a ratio as low as 1:10,000.

Although scFv and monoclonal antibodies (mAbs) targeting neoantigen P53^R175H^ have been reported^25, 26^, P53^R175H^-specific nanobodies have not yet been identified. Nanobodies are emerging as advanced antibody-derived tools for cancer diagnosis and therapeutics, offering better solubility, stability, and expression efficiency than scFvs, with strong potential for intracellular targeting and modular design^27, 28^. We aimed to determine whether the E-A functional screening strategy could identify nanobodies specific to neoantigen P53^R175H^ presented by HLA-A*02:01. We first generated a naïve alpaca VHH library with an estimated diversity of >10^13^, and employed phage display method to reduce the library size to 10^8^ through bio-panning by HLA-A*02:01-P53 R175H mut monomer and WT monomer (**Supplementary Fig. 7b**, c).

Then the pre-enriched VHH library was constructed into STAR-JC5 cells as a VHH lib STAR-JC5 cells library and performed E-A functional screening (**Fig 5d and Supplementary Fig. 7d**, e). In the first round of VHH STAR-JC5 cell screening, we utilized T2 cells loaded with the P53 R175H mut peptides as target cells for co-culture with the JC5 cell library as positive selection. We found that the percentage of E-A functional index-positive cells reached 20%, and the TCR^-^CD69^+^ cells were sorted for a second round of screening using the WT peptide-loaded T2 cells, referred to as negative selection, with the aim of eliminating non-specific binding and self-activating cells. To further enrich the positive cells, another positive selection was performed as third round of screening, resulting in an increased E-A functional index-positive rate of 32.1%. The positive JC5 cells sorted from the third round of functional screening were subjected to NGS sequencing to identify the selected nanobodies (**Supplementary Fig. 7f**). Notably, top five VHHs sequences showed enrichment (**Fig. 5e**). We transduced human primary T cells with the top five enriched VHHs-based mutSTAR (the surface display levels of mutSTAR receptors on T cells were shown in **Supplementary Fig. 8a**) and co-cultured them with T2 cells loaded with P53 R175H mut peptide or wt peptide. Consistently, the top three P53 VHH mutSTAR-T cells exhibited dose-dependent activation upon stimulation with serial dilutions of P53 R175H peptides, with a half maximal effective concentration (EC_50_) of 1.603 nM for P53-1 VHH, 1.482 nM for P53-2 VHH and 10.87nM for P53-3 VHH (**Fig. 5f**). In contrast, no responses were observed for the last two VHH groups.

### Novel antibody-based STAR-T cells exhibit potent anti-tumor efficacy

To assess the potential of clinical application for the top three functional VHHs, the cytotoxicity and cytokines release (IFNγ, TNFα and IL2) of the three P53 VHHs-based mutSTAR-T cells were further evaluated (**Fig. 6a**). Bioluminescence-based cytotoxicity assay revealed substantial killing of T2 cells loaded with P53 mut peptides for these three STAR-T cells, while the P53-2 VHH mutSTAR-T cells exhibited a minimal level of non-specific cell lysis under high concentration of wt peptides. And these three STAR-T cell groups showed substantial cytokines secretion upon P53 mut peptides stimulation. Taken together, these data indicate that three of the top five enriched VHHs discovered by E-A functional screening could recognize neoantigen P53^R175H^ in the context of HLA-A*02:01, and these VHHs-based STAR-T cells effectively respond to eliminate target tumor cells *in vitro*.

**Fig. 6.**
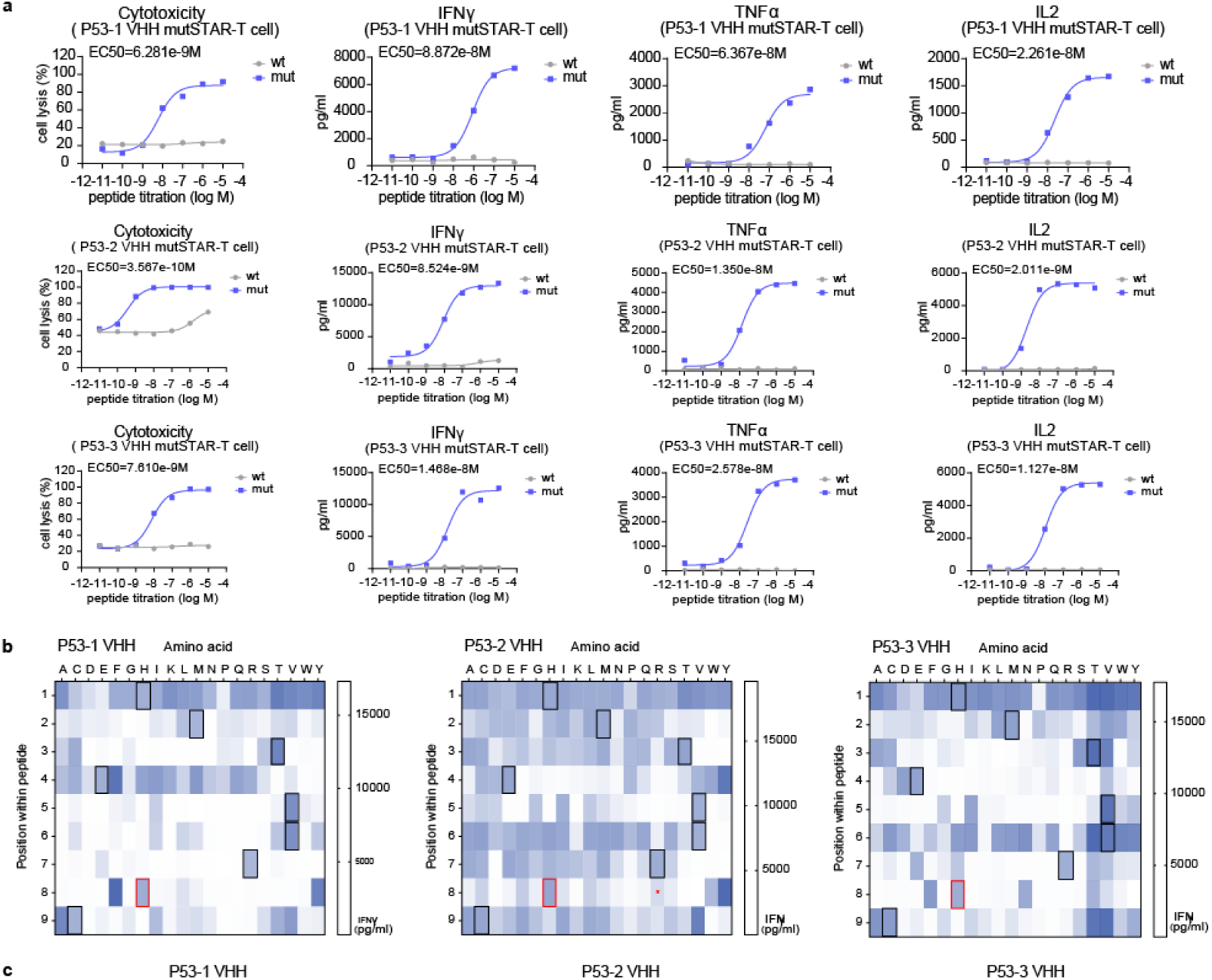
Neoantigen P53^R175H^ specific tumor responsiveness *in vitro* and identification of putative cross-reactive peptides. **a**, Cytotoxicity (presented as percentage of specific cell lysis) and cytokine release (IFN-γ, TNF-α, IL-2) of top three-ranked E-A functional index-positive P53 VHHs-based mutSTAR-T cells against T2 cells loaded with different doses of P53 wt peptide or R175H mut peptide. **b, c**, Peptide scanning mutagenesis to assess potential P53-1, P53-2 and P53-3 VHH cross-reactivity. Each amino acid of the P53 R175H peptides was systematically substituted with the other common 19 amino acids, thereby generating libraries of variant peptides each differing from the original peptide by a single amino acid. T2 cells were pulsed with 10 μM of individual peptides from the R175H peptide library, and were combined with P53 VHHs mutSTAR-T cells. **b,** Supernatant was assayed for IFN-γ, with the mean of three technical replicates plotted as a heatmap. Black boxes indicate amino acids in the parental peptides. Red boxes indicate the mutation “H” site. Red asterisks indicate the wt “R” site. **c**, Illustration of the binding patterns of P53 VHHs as Seq2Logo graphs, calculated by dividing the IFN-γ value by 10^5^ and using the PSSM-Logo algorithm. The co-culture time used in the experiments is 24h and the E:T ratio is 1:1. EC50, half maximal effective concentration. Data in (**a**) are representative of three independent experiments.

One of the major challenges in developing novel immunotherapeutic antibodies is off-target binding, which may induce toxicity in normal cells. To assess the cross-reaction of the top three P53 VHHs, we performed scanning mutagenesis to identify peptides in the human proteome. A peptide library was constructed by systemically substituting amino acids at each position of the target P53 R175H peptide with each of the remaining 19 common amino acids. T2 cells loaded with each of the 171 variant peptides were then used to assess T cell activation by measuring IFN-γ release after co-cultured with P53 VHHs mutSTAR-T cells (**Fig. 6b**). Compared to P53-2 VHH, both P53-1 VHH and P53-3 VHH exhibited greater specificity in recognizing the peptide library, particularly at positions located in the C-terminal half of the peptide where the mutant residue resides. Most changes at these positions abolished VHH recognition. In contrast, substitutions at the N-terminal positions were often tolerated without substantially affecting the interaction between the VHHs and the peptides. This recognition pattern was illustrated as a Seq2Logo graph (**Fig. 6c**)

We further evaluated the binding affinity using SPR, which revealed that the P53-1,2,3 VHHs bound to P53 R175H-HLA-A*02:01 monomer with a dissociation constant (Kd) of 2.85 μM, 375 nM, 5.25 μM, respectively (**Supplementary Fig. 8b**).

The above analyses reveal that the P53-1 VHH was highly enriched and demonstrated proper immune responsiveness against tumor cells *in vitro*. Therefore, the P53-1 VHH-based mutSTAR-T cells were selected for further evaluation of *in vivo* anti-tumor efficacy. We engrafted KMS26 multiple myeloma cells which endogenously express P53^R175H^ neoantigen and HLA-A*02:01 into NCG mice through intravenous injection, establishing widespread, actively growing cancers throughout the body (**Fig. 7a**). The mice were divided into two groups according to luminescence quantification of tumor burden at 4h after tumor inoculation, and the P53-1 VHH mutSTAR-T cells were infused intravenously at day 3. Mock-T cells with exogenous RFP transduction were infused as negative controls. The tumor burden were monitored every few days. We found that the tumor growth was markedly suppressed in the P53-1 VHH group (P<0.0001) (**Fig. 7b, c**). Meanwhile, we also established an over-expressed K562 mouse model which stably expressing the P53^R175H^ tandem minigenes (K562-A2-GFPM3-luc) or P53 wt protein (K562-A2-P53 wt-luc) (**Fig. 7d, g**). We subcutaneously engrafted K562 tumor cells into NCG mice respectively, and divided the mice into two groups according to luminescence quantification of tumor burden. The P53-1 VHH mutSTAR-T cells were infused intravenously at day 3. Mock-T cells with exogenous RFP transduction were infused as negative controls and the tumor burden were monitored. We found that the tumors expressing P53 mut neoantigen were significantly suppressed in the P53-1 VHH group (P<0.0001), whereas tumors expressing P53 wt protein showed no noticeable change (**Fig. 7e, f, h, i**). These *in vivo* experiments indicate that the novel P53 VHH-based STAR-T cells exhibit potent anti-tumor efficacy, and demonstrate that the E-A functional screening enables the effective identification of neoantigen-specific antibodies with promising anti-tumor activity.

**Fig. 7.**
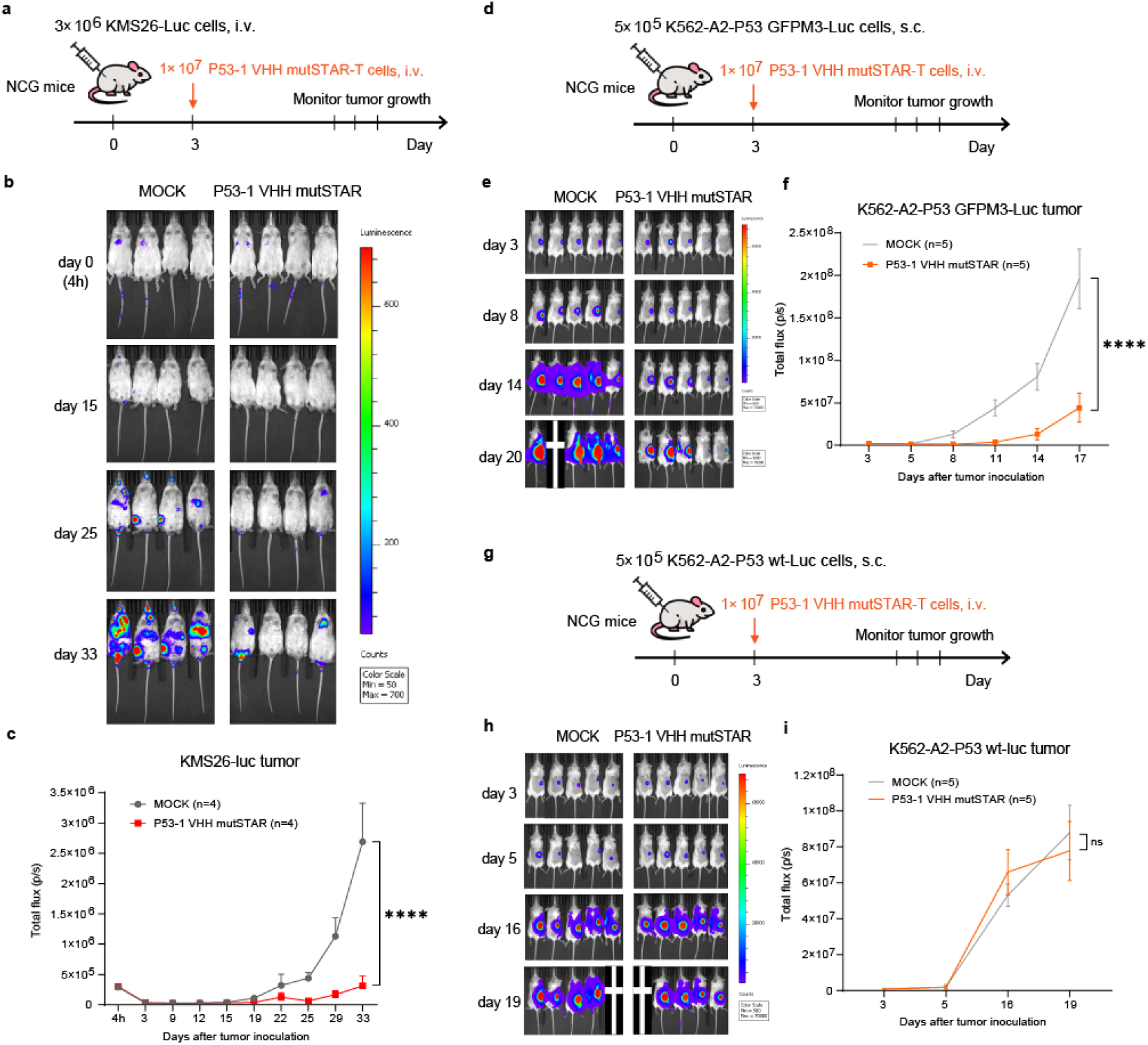
Tumor inhibition efficacy *in vivo* of P53-1 VHH mutSTAR-T cells. **a**, Schematic of the experimental process for KMS26 mouse model which endogenously express P53^R175H^ neoantigen and HLA-A*02:01. NCG mice were inoculated intravenously with 3×10^6^ KMS26-Luc cells at day 0, and 1×10^7^ P53-1 VHH mutSTAR-T cells were intravenously injected on day 3. Tumor growth was monitored by *in vivo* luminescence imaging. **b**, Bioluminescence images of KMS26-Luc tumor, with 4 mice in each group and MOCK group were infused with T cells expressing RFP. **c**, KMS26-Luc tumor progression as evaluated by bioluminescence imaging. **d, g**, Schematic of the experimental process for K562 over-expression mouse model. NCG mice were inoculated subcutaneously with 5×10^5^ K562-A2-P53 GFPM3-Luc cells stably expressing P53^R175H^ neoantigen or K562-A2-P53 wt-Luc cells stably expressing P53 wt protein at day 0, and 1×10^7^ P53-1 VHH mutSTAR-T cells were intravenously injected on day 3. Tumor growth was monitored by *in vivo* luminescence imaging. **e, h**, Bioluminescence images of K562 tumor, with 5 mice in each group and MOCK group were infused with T cells expressing RFP. **f, i**, K562 tumor progression as evaluated by bioluminescence imaging. Statistical analysis of quantification in (**c, f, i**) was performed using two-way analysis of variance (ANOVA). Data are representative of three independent experiments and are presented as mean ± SEM. P values were presented as indicated. ****P < 0.0001, ns means no significant difference (P>0.05).

### Screened novel antibodies can be adapted into multiple therapeutic formats

Given the diverse T cell-based approaches available for cancer-targeted therapies, we further converted the screened P53-1 VHH into CARs format or Bi-specific antibody (BsAb) format to assess its anti-tumor functionality. We first constructed P53-1 VHH CAR with either 4-1BB (BBzCAR) or CD28 (28zCAR) costimulatory domains as previously described^5^, and transduced into T cells to validate their functions (**Fig. 8a, b**). After co-incubated them with T2 cells loaded with P53 R175H mut peptide or wt peptide, the P53-1 VHH CAR-T cells exhibited similar antigen-specific activation, tumor cell killing ability and IFNγ release. At the meantime, we produced P53-1 VHH BsAb including anti-CD3 scFv and Fc domain, which can simultaneously bind to tumor cells via tumor-specific antigen P53^R175H^ and to T cells through surface CD3 (**Fig. 8c**). To assess an applicable concentration of target antigen required for activating T cells and killing tumor cells *in vitro*, we pulsed T2 cells with 10^-4^M ∼10^-7^M P53 R175H mut peptide or wt peptide and co-cultured them with T cells in the presence of 10nM P53-1 VHH BsAb. We found the T cells exhibit adequate CD69 activation level and tumor cell killing ability with 10^-4^M R175H peptide concentration. To determine the minimal effective dose, we reduced the concentration of BsAb in co-culture from 10nM to 0.1nM and found that the T cell function exhibit BsAb-dose dependent with 1nM of BsAb capable of triggering distinct activation and cytotoxicity of T cells.

**Fig. 8.**
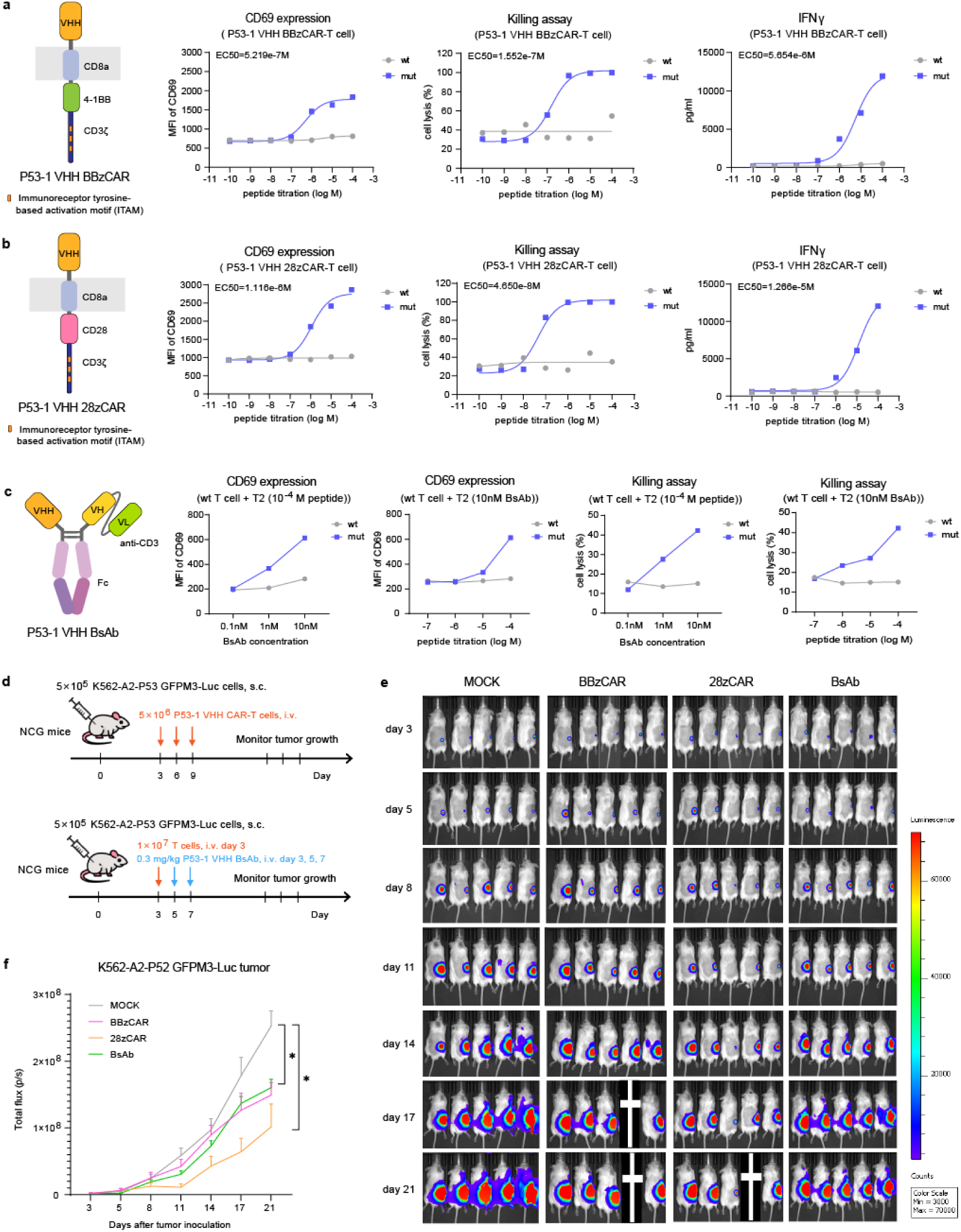
CAR-T and BsAb armed with nanobodies from E-A functional screening exhibit potent anti-tumor efficacy both *in vivo* and *in vitro*. **a**, Schematic structure of P53-1 VHH BBzCAR and the *in vitro* anti-tumor functions (including CD69 activation, killing ability, IFN-γ secretion) of P53-1 VHH BBzCAR-T cells against T2 cells loaded with different doses of P53 wt peptide or R175H mut peptide. **b**, Schematic structure of P53-1 VHH 28zCAR and the *in vitro* anti-tumor functions of P53-1 VHH 28zCAR-T cells. **c**, Schematic structure and the *in vitro* anti-tumor functions of P53-1 VHH BsAb. The co-culture time used in the experiments **a**-**c** is 20h and the E:T ratio is 1:1. EC50, half maximal effective concentration. **d**, Schematic of the experimental process for K562-A2-P53 GFPM3-Luc mouse model treated by P53-1 VHH BBzCAR-T cells, P53-1 VHH 28zCAR-T cells and P53-1 VHH BsAb. NCG mice were inoculated subcutaneously with K562-A2-P53 GFPM3-Luc cells (5×10^5^) stably expressing P53^R175H^ neoantigen at day 0. For CAR-T groups, three dose of P53-1 VHH CAR-T cells (5×10^6^) were intravenously infused on day 3, 6 and 9. For BsAb group, human T cells (1×10^7^) were intravenously infused on day 3, and three doses of P53-1 VHH BsAb at 0.3 mg/kg were administrated through intraperitoneal injection at day 3, 5 and 7. Tumor growth was monitored by *in vivo* luminescence imaging. **e**, Bioluminescence images of K562-A2-P53 GFPM3-Luc tumor with 5 mice in each group. **f**, K562-A2-P53 GFPM3-Luc tumor progression as evaluated by bioluminescence imaging. Statistical analysis of quantification in (**f**) was performed using two-way analysis of variance (ANOVA). Data are representative of three independent experiments and are presented as mean ± SEM. P values were presented as indicated. *P < 0.1.

Furthermore, comparable *in vivo* determinations were executed for the P53-1 VHH CAR-T and BsAb. We first subcutaneously engrafted K562-A2-P53 GFPM3-Luc cells in NCG mouse model, and the mice were divided into two groups according to luminescence quantification of tumor burden at day 3 after engraftment. For CAR groups, the P53-1 VHH BBzCAR-T cells or P53-1 VHH 28zCAR-T cells were infused intravenously at day 3, 6 and 9 (**Fig. 8d**). For BsAb group, mice were inoculated 1×10^7^ *in vitro* expanded human T cells via lateral tail vein injection on day 3, and subsequently received intraperitoneal injection of P53-1 VHH BsAb at day 3, 5 and 7 (**Fig. 8d**). Mock human T cells with no transduction were infused as negative controls. The tumor burden were monitored every few days (**Fig. 8e**). We found that the tumor growth was significantly suppressed in the CAR and BsAb groups (P<0.1), which indicated that the novel P53 VHH-based multiple T cell therapies exhibit potent anti-tumor efficacy (**Fig. 8f**).

## Discussion

Over the past few decades, antibody therapeutics in the treatment of cancer has witnessed remarkable advancements, resulting in a significant surge in the demand for high-affinity and specific antibodies^29^. Traditional antibody discovery methods, such as phage display and hybridoma technologies, have been instrumental in identifying antibodies against tumor-associated antigens (TAAs). However, these approaches often require extensive functional validation to confirm the biological activity of the antibodies. phage display methods are also restricted by non-specific binding of phage to reagents or consumables and subsequent propagation throughout each iteration^30^. In contrast, our study introduces a novel functional screening strategy, termed E-A (Endocytosis-Activation) functional screening, which leverages the intrinsic properties of T cells to directly identify functional antibodies, particularly those targeting MHC-presented neoantigens.

Traditional cell panning techniques, such as those described by Alonso-Camino et al.^31^, involve the use of chimeric antigen receptors (CARs) to display antibody libraries on the surface of T cells. These CAR-T cells are then subjected to multiple rounds of selection to enrich for antibodies with desired specificities. While effective, these methods can be limited by issues such as tonic signaling and auto-activation of T cells in the absence of antigen, which may confound the identification of functional antibodies. Our E-A functional screening approach addresses these limitations by utilizing two distinct functional readouts: STAR receptor endocytosis and T cell activation (CD69 upregulation). This dual-readout system ensures that only antibodies capable of both antigen engagement and subsequent T cell activation are selected, thereby increasing the likelihood of identifying truly functional antibodies. These indicators have been proven to exhibit strict antigen specificity and sensitivity with low signal-to-noise ratio. In contrast, although CAR also has the capability of antibody carrying, it triggers noticeable tonic signaling and T cell auto-activation under antigen-free conditions, which limits them for functional screening^5^. Furthermore, by directly assessing functional activity, our method reduces the need for extensive post-selection validation, streamlining the antibody discovery process. Notably, considering cell culture conditions and infection efficiency, the size of the cell library utilized for the functional screening method may be limited to not exceeding 10^9^. Rigorous positive and negative selection is necessary for our E-A functional screening, otherwise, non-specific or non-functional clones may confound the screening results.

Furthermore, E-A functional screening not only applies to screening antibody targeting tumor-associated antigens (TAAs) but also facilitates the discovery of TCR-like antibody targeting tumor-specific neoantigen. Targeting TAAs has been reported to lead to on-target/off-tumor toxicity and resistance, due to TAA expression in normal tissues and the loss of TAA expression in tumor cells^32^. These drawbacks can be circumvented by targeting neoantigens, which offer a distinct advantage with their unique tumor specificity and absence in normal tissues, making them ideal targets for personalized tumor treatment^33, 34^. Regarding TCR-like antibodies, these are a novel class of antibodies that can recognize peptide/MHC complexes on tumor cell surfaces, mimicking the specificity of T cell receptors, which have shown promise in cancer immunotherapy by targeting intracellular tumor-specific antigens presented on the cell surface^35^. These TCR-like antibodies can easily be transformed into various therapeutic formats, including full-length antibodies, ADCs, BsAbs, CAR and STAR. CAR-T cells with an scFv that recognizes the oncogene nucleophosmin (NPM1c) epitope-HLA-A2 complex demonstrate strong cytotoxicity against NPM1c^+^HLA-A2^+^ leukemia cells and acute myelocytic leukemia (AML) blasts with minimal to on-target/off-tumor toxicity^36^. The single-chain diabody (scDb) with an scFv specific to the HLA-A3-restricted KRAS neoantigen effectively activate T cells to lyse target cancer cells expressing endogenous levels of the mutant RAS proteins and cognate HLA alleles^37^. Our recent work reported that TCR-mimicking STAR, which combines TCR-like antibody, exhibited high antigen sensitivity when targeting low-density neoantigen^38^. These neoantigen-specific antibodies are typically screened using phage or yeast surface display methods combined with synthetic or immunized libraries, which are affinity-based and require additional functional verification. However, our E-A functional screening strategy enables the identification of multiple novel neoantigen-specific antibodies with certain anti-tumor functions and boasts a high screening positive rate, thus saving time and costs. More importantly, the screened antibodies can be converted into multiple therapeutic antibody formats, each demonstrating effective anti-tumor efficacy.

In our study, we utilized the Endocytosis-Activation (E-A) screening platform to identify nanobodies, specifically minibinders, that target neoantigens presented by HLA molecules on tumor cells. A key feature of our work is that the nanobodies, which are compact, single-domain minibinders, retain high affinity and specificity. These minibinders are much smaller than conventional antibodies, making them ideal candidates for targeted cancer immunotherapy. By using the E-A screening strategy, we were able to efficiently isolate minibinders that specifically recognize neoantigens, such as p53 mutations presented by HLA molecules on tumor cells. This marks a significant step forward, as minibinders—unlike conventional antibodies—are not only smaller but can also be engineered to bind with higher precision, providing more efficient tumor targeting. The small size of these minibinders also translates into better tumor penetration and potentially lower immunogenicity, thus representing a promising direction for the development of personalized cancer therapies.

In summary, we have developed an efficient E-A functional antibody screening strategy for targeted cancer therapy. Our study demonstrated the applicability of E-A functional screening in identifying both TAA- and neoantigen-specific nanobodies. Additionally, other antibody formats, such as scFv, could also be utilized in this screening system. Further study is needed to optimize the antibodies screened for superior functionality and assess their potential clinical application. Nevertheless, E-A functional screening holds considerable value in adoptive T cell therapy for cancer treatment by providing a highly specific and efficient method for identifying antibodies that target tumor-specific MHC-presented neoantigens, thereby improving the safety, efficacy, and personalized nature of cancer immunotherapy.

## Methods

### Cell lines and primary cells

Lenti-X 293T cells were purchased from Takara Biomedical Technology Co., Ltd. KMS26, K562, Raji and Jurkat E6.1 cells were purchased from American Type Culture Collection (ATCC). JC5 cells were derived from Jurkat E6.1 cells by knocking out TCRα and TCRβ chains with a CRISPR-Cas9 system [guide RNA (gRNA) sequences: TRBC_ GGGCTCAAACACAGCGACCTC and TRAC_GTCTCTCAGCTGGTACACGGC]. T2 cells were obtained from BriStar Immunotech Co., Ltd. Primary human peripheral blood mononuclear cells (PBMCs) were purchased from Shanghai Oribiotech Biotechnology Co., Ltd. Lenti-X 293T cells were cultured in DMEM (Gibco, cat.C11995500BT) supplemented with 10% (v/v) FBS (ExCell Bio, cat.12A056) and 100 IU/ml penicillin-streptomycin (Yeasen, cat.60162ES76). KMS26, K562, Raji, Jurkat E6.1, JC5 and T2 cells were cultured in RPMI 1640 (Gibco, cat.C11875500BT) supplemented with 10% (v/v) FBS (ExCell Bio, cat.12A056) and 100 IU/ml penicillin-streptomycin (Yeasen, cat.60162ES76). Primary human PBMCs were cultured in RPMI 1640 (Invitrogen, cat.11875135) supplemented with 10% (v/v) heat-inactivated FBS (Gibco, cat.10091148), 100 IU/ml penicillin-streptomycin (Yeasen, cat.60162ES76) and 200 IU/mL rhIL-2 (Peprotech, cat.200-02). All cells were cultured at 37 °C with 5% atmospheric CO_2_.

### Vector construction

Lentiviral vectors (pHAGE) were used for transducing K562 target cell line which plasmid backbone contained an EF1α promoter and an IRES-linker GFP reporter. A fusion protein gene (P53 GFPM3) of GFP full length gene and P53 R175H (<u>HMTEVVR**H**C</u>) tandem minigenes (3×TACAAGCAGTCACAG<u>CACATGACGGAGGTTGTGAGGCACTGC</u>CCCCAC CATGAGCGC, coding protein: 3×YKQSQ<u>HMTEVVR**H**C</u>PHHER) was constructed in a pHAGE plasmid for transducing *in vivo* K562-A2-P53 GFPM3 tumor cell line. Full length P53 wt gene was constructed in a pHAGE plasmid for transducing *in vivo* K562-A2-P53 wt tumor cell line. Luciferase-encoding genes were constructed in pHAGE plasmids carrying puromycin resistance gene as an effective selection marker or GFP reporter gene. Another pHAGE vector which plasmid backbone contained an hEF1α-HTLV promoter and an IRES-linker RFP reporter were used for transducing T cell lines and primary T cells. The plasmids backbone structures of STAR, mutSTAR, BBzCAR and 28zCAR were as previously described^5^. In brief, STAR was designed as an antibody-TCR chimera by ligating human TCR constant regions with variable regions of antibody, and mutSTAR was a mutant of STAR by replacing human TCR constant regions with murine counterparts and introducing an additional interchain disulfide bond within TCRα/β constant regions and hydrophobic substitutions to TCRα transmembrane domain. CD3ζ cytoplasmic domain (UniProtKB-P20963, amino acids 52 to 164), 4-1BB cytoplasmic domain (UniProtKB-Q07011, amino acids 214 to 255) and CD28 cytoplasmic domain (UniProtKB-P10747, amino acids 180 to 220). Sequences for VHHs and scFvs used in this study were as previously described: MSLN VHH (PCT/CN2022/130106), CD123 VHH (US20160251440), CD19 scFv (FMC63)^39^, GPC3 scFv (GC33)^40^, CD3 scFv (OKT3)^41^. Gene fragments were assembled by seamless cloning kit (Clone Smarter, #C5891-50) into vector backbones together with a Kozak sequence for optimal translation.

### Lentivirus production

Lentiviruses encoding MSLN, CD123, CD19, GPC3, CD22, P53 GFPM3, P53 wt, HLA-A*02:01 and luciferase were packaged using packing and envelope plasmids psPAX2 and pMD2.G in Lenti-X 293T cells. Lentiviruses encoding VHH STARs, VHH STAR libraries, VHH mutSTARs and VHH CARs were packaged using packing and envelope plasmids pMD2.G, pMDLg/pRRE and pRSV-Rev in Lenti-X 293T cells. Plasmids were transfected into cells using polyethyleneimine (Polysciences Inc., #24765-1). The virus-containing culture medium supernatants were collected at 48 and 72 hours after transfection and filtered through a 0.45-μm syringe filter. The viruses were concentrated by mixing with PEG8000 (Sigma-Aldrich) overnight at 4°C and centrifuged for 30 min at 3,500 rpm.

### Cell lines construction

KMS26, K562, Raji and T2 cells were engineered and used as target cells. JC5 cells and primary human T cells were engineered and used as effector cells. KMS26, K562, Raji, T2 and JC5 cells were spin-infected with concentrated viral supernatants supplemented with 10 μg/ml Polybrene (Yeasen, cat. 40804ES86) for 90 min at 1,500 rpm at 32 °C and replaced with fresh RPMI 1640 medium 12 hours later. On day 3 post-infection, the reporter-positive target cells were sorted by FACS (fluorescence-activated cell sorting) or puromycin-resistant cells were selected to establish derivative cell lines as indicated. K562 cells stably expressing HLA-A*02:01 (K562-A2 cells) were prepared through transfection with lentiviruses of HLA-A*02:01. Raji-CD22KO cells were derived from Raji cells by knocking out CD22 with a CRISPR-Cas9 system [gRNA sequence: GGTATCCGATCCAATTGCAG]. All target cell lines were stably expressing luciferase through transfection with lentiviruses of luciferase.

### T cell activation and transduction

To transduce primary human T cells, PBMCs (1×10^6^ cells/ml) were activated in 48-well plates coated with 5μg/ml anti-human CD3 (Biolegend, cat.317326), 1μg/ml anti-human CD28 (BD Biosciences, cat.555725) and 5μg/ml human fibronectin (BD Biosciences, cat.354008) in T cell medium (PBMCs medium). After 36 hours of activation, the cells were spin-infected with concentrated viral supernatants for 90 min at 1,700 rpm at 32 °C, and replaced with fresh T cell medium 24 hours later. The transduction efficiency was determined using flow cytometry 72 hours after primary T cells were transduced.

### Library generation

All animal experiments were approved by the Committee on the Ethics of Animal Experiments of the Institute of Microbiology, Chinese Academy of Science (IMCAS) and conducted in compliance with the recommendations in the Guide for the Care and Use of Laboratory Animals of IMCAS Ethics Committee.

The CD22 immune VHH library was obtained from BriStar Immunotech Co.,Ltd. by immunization of alpaca. In brief, a healthy alpaca was immunized subcutaneously in the neck area with 200 μg of CD22 protein mixed with MnJ adjuvant every two weeks. A week after the sixth immunization, peripheral blood of the alpaca was collected and B cells were sorted using B Cell Isolation Kit (Stemcell, cat. 17954_C). RNA of the sorted B cells was extracted using FastPure Cell/Tissue Total RNA Isolation Kit (Vazyme, cat. RC112-01) and reverse transcribed into cDNA which was used as template in PCR to obtain VHH library with the following primers: VHH-Forward, 5’-GATGTGCAGCTGCAGGAGTC-3’; VHH-Reverse, 5’-TGAGGAGACGGTGACCTGGGT-3’. The VHH library was then constructed to STAR pHAGE plasmid under the control of hEF1α-HTLV promoter and GM-CSF signal peptide (before TCRBC chain). Then the CD22 VHH STAR library was transduced to JC5 cells by lentiviruses infection to generate CD22 VHH lib STAR-JC5 cell library.

Naïve VHH library was obtained from BriStar Immunotech Co., Ltd. by collection of healthy (never challenged by a specific antigen) alpacas’ peripheral bloods. In brief, peripheral blood of healthy alpaca was collected and total B cells were sorted using B Cell Isolation Kit (Stemcell, cat. 17954_C). RNA of the sorted B cells was extracted using FastPure Cell/Tissue Total RNA Isolation Kit (Vazyme, cat. RC112-01) and reverse transcribed into cDNA which was used as template in PCR to obtain VHH library with VHH-Forward and VHH-Reverse primers. The VHH library was then constructed to pADL-22c phagemid vector to obtained naïve VHH phage display library. After four rounds of phage bio-panning, the pre-enriched VHH library was cloned and constructed to STAR pHAGE plasmid under the control of hEF1α-HTLV promoter and GM-CSF signal peptide (before TCRBC chain). Then the VHH STAR library was transduced to JC5 cells by lentiviruses infection to generate pre-enriched VHH lib STAR-JC5 cell library.

### Peptide loading

The peptides in this study were all synthesized by Nanjing GenScript Biotech Corporation at a purity of >98%. Peptides were dissolved in DMSO at 10 mM working stock and stored at −20°C. Gradient dilutions of the peptides were performed using RPMI 1640 medium. Target cells (5×10^5^ cells/ml) were loaded with specified concentration of peptides for 2 hours at 37°C. After incubation, the cells were washed three times with PBS and resuspended with medium for co-incubation experiments.

### BsAb production

Using the heterodimeric Fc variant technology KiHss-AkKh, as previously described^42^, we fused the VHH fragment of P53-1 VHH to the Fc-engineered variant (LALAPG), and the anti-CD3 scFv to the hole variant Fc region. The P53-1 VHH bispecific antibody (BsAb) was produced via transient co-transfection of plasmids encoding both arms into FreeStyle 293-F cells. The supernatant containing BsAbs was purified using Protein A affinity chromatography, following the manufacturer’s instructions. The purified proteins were desalted into PBS using 50 kDa ultrafiltration discs and subsequently filtered through a 0.22 µm membrane to remove aggregates and achieve sterility. The heterogeneity and purity of the proteins were then confirmed by SDS-PAGE analysis.

### Flow cytometry

Cells were stained with fluorophore-conjugated antibodies (0.5μg/ml) in PBS at a concentration of 1×10^7^ cells/ml. The following antibodies were used: anti-human TCRα/β-BV421 (Biolegend, cat.306722), anti-human CD69-APC (Biolegend, cat.310910), anti-mouse TCRβ-APC (Biolegend, cat.109212), anti-human CD69-APC/Cy7 (Biolegend, cat.310914), fixable viability dye eFluor 506 (eBioscience, #65-0866-18) and HLA-A2-APC (Biolegend, cat. 343307). Flow cytometry data were acquired by BD Fortessa and analyzed by using FlowJo software (BD Biosciences).

### Cytotoxicity assay

In order to establish target cells for bioluminescence-based cytotoxicity assay, a panel of cell lines were generated via lentiviral transduction including Raji-luciferase, Raji-CD22KO-luciferase, KMS26-luciferase, K562-A2-P53 GFPM3-luciferase, K562-A2-P53 wt-luciferase and T2-luciferase. 100,000 target cells in T cell medium were seed in 96-well-plates co-cultured with effector T cells at 1:1 E: T ratios for 24 hours in triplicate. Wells containing only target cells served as negative controls for baseline cytotoxicity. Subsequently, the supernatant was removed and the collected cells were treated with 100 μL 1 × RIPA lysis buffer (Beyotime, cat. P0013B) on ice for 30 min. After centrifuged for 10 min at 3,500 rpm, 50 µL lysate supernatant were mixed with 50 µL Luciferase reagent (Yeasen, cat.11401ES80) in a LumaPlate-96 white plate (Nunc, cat.463201), followed by measuring BLI with a luminometer (Promega GloMax® 96 Microplate Luminometer). The % specific lysis of tumor cells was calculated using the formula: % specific lysis = (luminescence of target cells - luminescence of target cells cocultured with effector cells) / luminescence of target cells × 100.

### Enzyme-linked immunosorbent assay

The cytokines responses of T cells were determined by cytokines released into culture medium. The cytokines concentrations were measured using the following enzyme-linked immunosorbent assay kits per the manufacturer’s instructions: human IFN-γ (Invitrogen, cat.88-7316-88), human TNF-α (eBioscience, cat.88-7346-88) and human IL-2 (Invitrogen, cat.88-7025-88).

### *In vivo* anti-tumor activity in a xenograft tumor model

Six- to eight-week-old male NCG mice were purchased from GemPharmatech Co. and kept under specific pathogen-free conditions in animal facility of the Tsinghua University. All mouse experiments were conducted according to Institutional Animal Care and Use Committee (IACUC)-approved protocols. Tumor burden was measured using a sensitive *in vivo* luminescence imaging (IVIS Spectrum, PerkinElmer) method after intraperitoneal D-Luciferin, Sodium Salt D (Yeasen, cat. 40902ES02) injection, and total flux was computed using the provided software (Living Image, Perkin Elmer), which measures the brightness through typical circular regions of interest (ROIs). According to their initial tumor burden, mice were serpentine randomized to mock and treatment groups. T cells were infused intravenously via tail vein in 100 μl volume. BsAb were injected intraperitoneally at 0.3 mg/kg in 200 μl volume per dose. Tumor progression was evaluated every 3 or 5 days, and the end points were determined by a tumor flux intensity above 5 × 10^8^. Mice were sacrificed with a CO_2_ chamber.

### PCR amplification and deep sequencing

Genomic DNA of sorted VHH STAR-JC5 cells was extracted using TIANamp Genomic DNA Kit (Tiangen, cat. DP304-02) and used as template in PCR amplification in a Bio-Red Thermal Cycler PCR System with the following program: initial denaturation step at 95 °C for 3 min followed by 25 cycles of 95 °C for 15 s, 59 °C for 15 s, and 70 °C for 30s, then a final extension at 70 °C for 3 min, followed by a 4°C hold. The primers used in PCR amplification were VHH-Forward and VHH-Reverse previously described. The amplified PCR products were purified using TIANgel Midi Purification Kit (Tiangen, cat. DP219-03) according to the manufacturer’s protocol. Sequencing was conducted by GENEWIZ using Illumina NovaSeq sequencing. Amino acid equences of VHHs identified in this study were listed in **supplementary table 1**.

### Statistical analysis

Statistical analyses were performed using GraphPad Prism 9 software. Statistical comparisons between two groups were determined by two-tailed Student’s *t*-tests for unpaired data. Statistical comparisons between multiple groups were determined by two-way analysis of variance (ANOVA). Data are reported as mean ± SEM. *P <0.05; **P < 0.01; ***P < 0.001; ****P < 0.0001; ns, not significant.

## Data availability

The original NGS DNA-seq data have been deposited in the Sequence Read Archive under accession numbers PRJNA1172294. The data that support the findings of this study are available from the corresponding author upon request.

## Acknowledgements

This work was partially supported by grants from National Key Research and Development Program of China (2019YFA0508502 and 2020YFA0509101 to X.L.), and grants from National Natural Science Foundation of China (31930039, 82293664, 31821003 to X.L.), and annual fund of Tsinghua-Peking Center for Life Sciences and Changping Laboratory.

## Author contributions

Y.L., D.H. and X.L. conceived of the approach and designed the research. Y.L., D.H., C.L. and Z.Z. performed experiments. Y.S. analyzed sequencing results. D.Y., W.R., Z.Y., L.Y. and J.W. provided critical reagents. Y.L., D.H., C.L., C.W., Z.Z., Y.F. and X.L. analyzed results. Y.L., D.H. and X.L. wrote the paper.

## Competing interests

X.L., Y.L. and D.H. are inventors on two patent applications (pending) concerning the described technology and the identified nanobodies in this research. The other authors declare no competing interests.

---

**Supplementary Figure 1.**
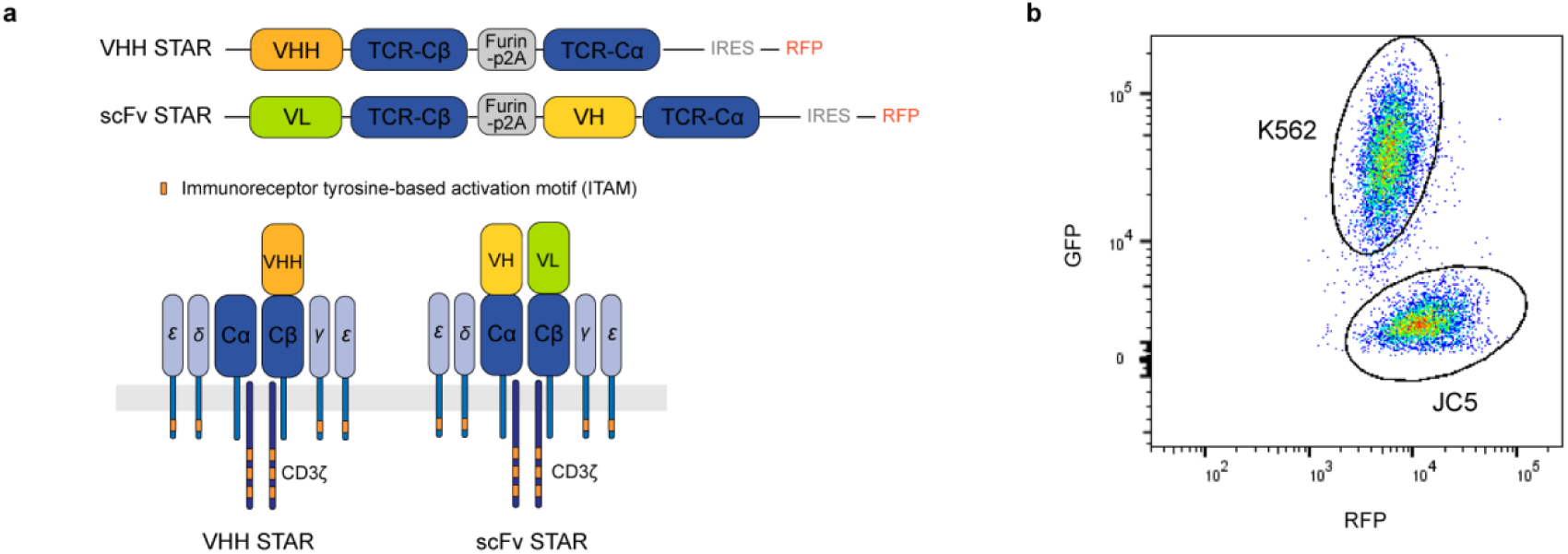
STAR constructs and gating strategy of FACS. (**a**) Schematic diagram and structure of VHH STAR and scFv STAR followed by an internal ribosome entry site (IRES) and a red fluorescent protein (RFP). (**b**) Resolution via flow cytometry of JC5 and K562 cells. The JC5 cells and K562 cells were resolved by gating on the RFP+ population and the GFP+ population, respectively.

**Supplementary Figure 2.**
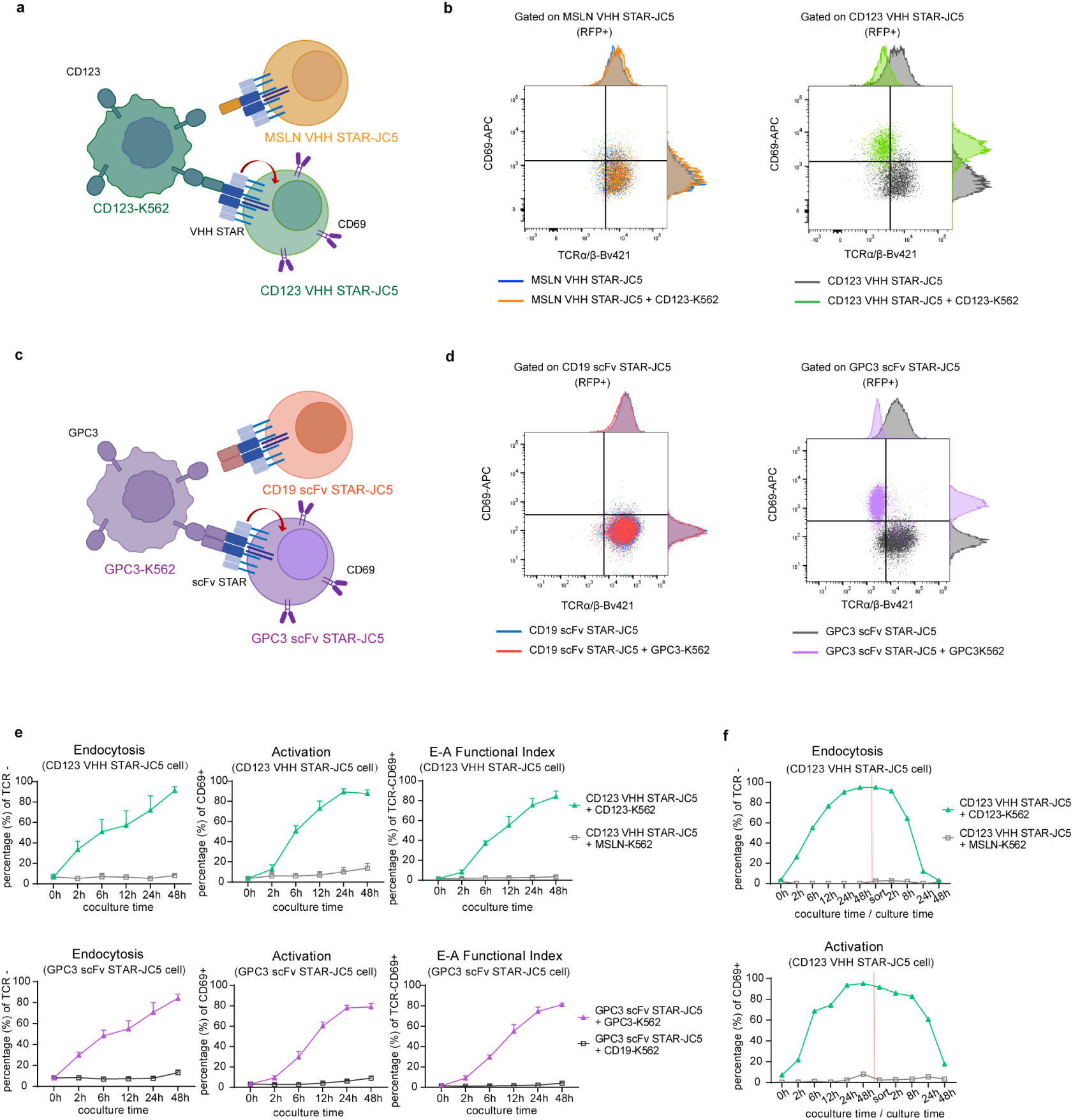
STAR endocytosis and CD69 activation of T cells occur in a membrane antigen-specific manner. (**a, c**) Schematic of the functional evaluation assay of the VHH STAR-JC5 cells and scFv STAR-JC5 cells. For VHH STAR group (**a**), K562 target cells were transduced to express tumor membrane antigen CD123 and subsequently co-cultured with JC5 cells transduced with cognate CD123 VHH STAR or non-cognate MSLN VHH STAR. For scFv STAR group (**c**), K562 target cells were transduced to express tumor membrane antigen GPC3 and subsequently co-cultured with JC5 cells transduced with cognate GPC3 scFv STAR or non-cognate CD19 scFv STAR. Same coloring indicates cognate antigen-antibody pairs. Red arrows represent endocytosis process. Purple homodimers represent surface molecular CD69. (**b, d**) The expressions of STAR and CD69 in RFP positive STAR-JC5 cells were detected by staining with antibodies specific to human TCRα/β and CD69 after co-culture for 24h. (**e**) Flow cytometry analysis of STAR endocytosis (TCRα/β negative), CD69 activation (CD69 positive) and E-A functional index (TCRα/β negative and CD69 positive) in RFP positive STAR-JC5 cells after co-incubation with target K562 cells expressing cognate or non-cognate membrane antigen at different time points. (**f**) Flow cytometry analysis of STAR-JC5 cells endocytosis and activation at different time point before and after JC5 cells were sorted at 48h of co-culture by FACS. The E:T ratio used in co-culture experiments is 1:1. Data in (**b**) and (**d-f**) are representative of three independent experiments. Data in (**e**) are presented as mean ± SEM.

**Supplementary Figure 3.**
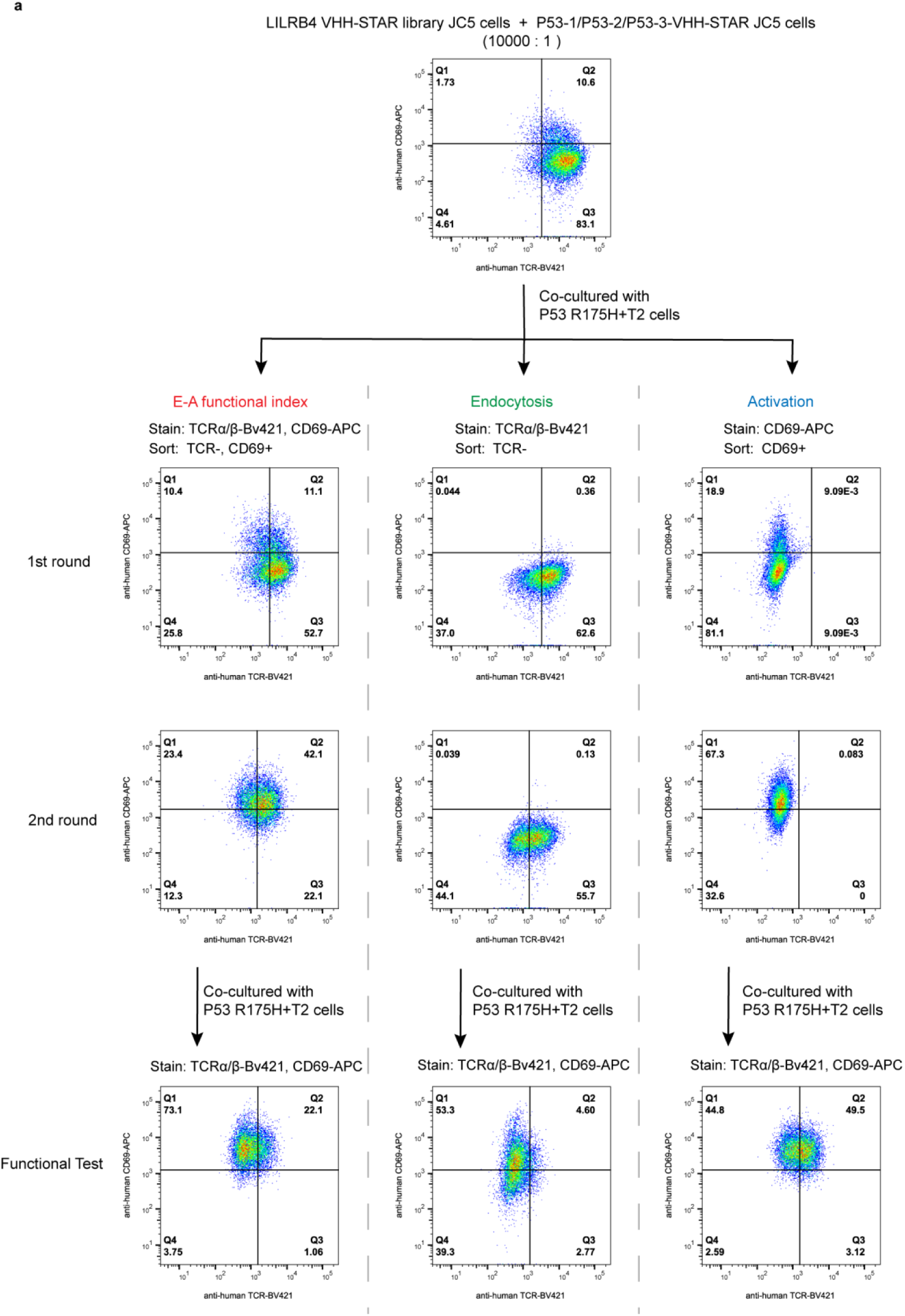
Comparative analysis of dual-readout E-A functional index (TCR endocytosis + CD69 upregulation), TCR-only, or CD69-only screening strategies. (**a**) A new LILRB4-specific VHH-STAR JC5 cell library, generated from an immunized alpaca and spiked with known functional clones (P53-1∼P53-3) at a 1:10,000 ratio, was co-cultured with P53 R175H loaded T2 target cells and subjected to two rounds of functional selection using either the dual-readout E-A functional index, TCR-only, or CD69-only strategies. Enriched populations were analyzed by flow cytometry for CD69 expression and TCR endocytosis following stimulation.

**Supplementary Figure 4.**
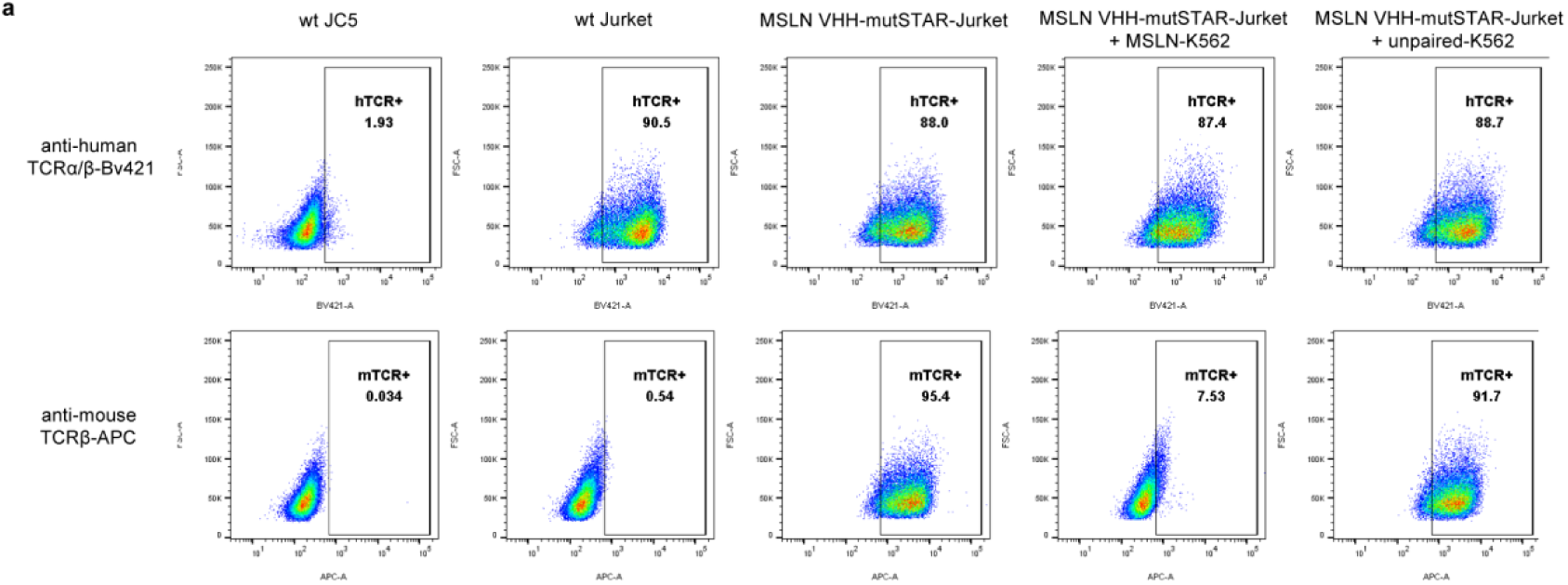
STAR endocytosis is directly linked to specific antigen binding. (**a**) TCR endocytosis rates of different T cells before and after co-incubate with target cells were assessed by flow cytometry. FSC were used to define the single cell population, the expression of human TCR was defined by staining with anti-human TCRα/β-Bv421 antibody and the expression of mutSTAR was defined by staining with anti-mouse TCRβ-APC antibody. The positive population was selected. Data are representative of three independent experiments.

**Supplementary Figure 5.**
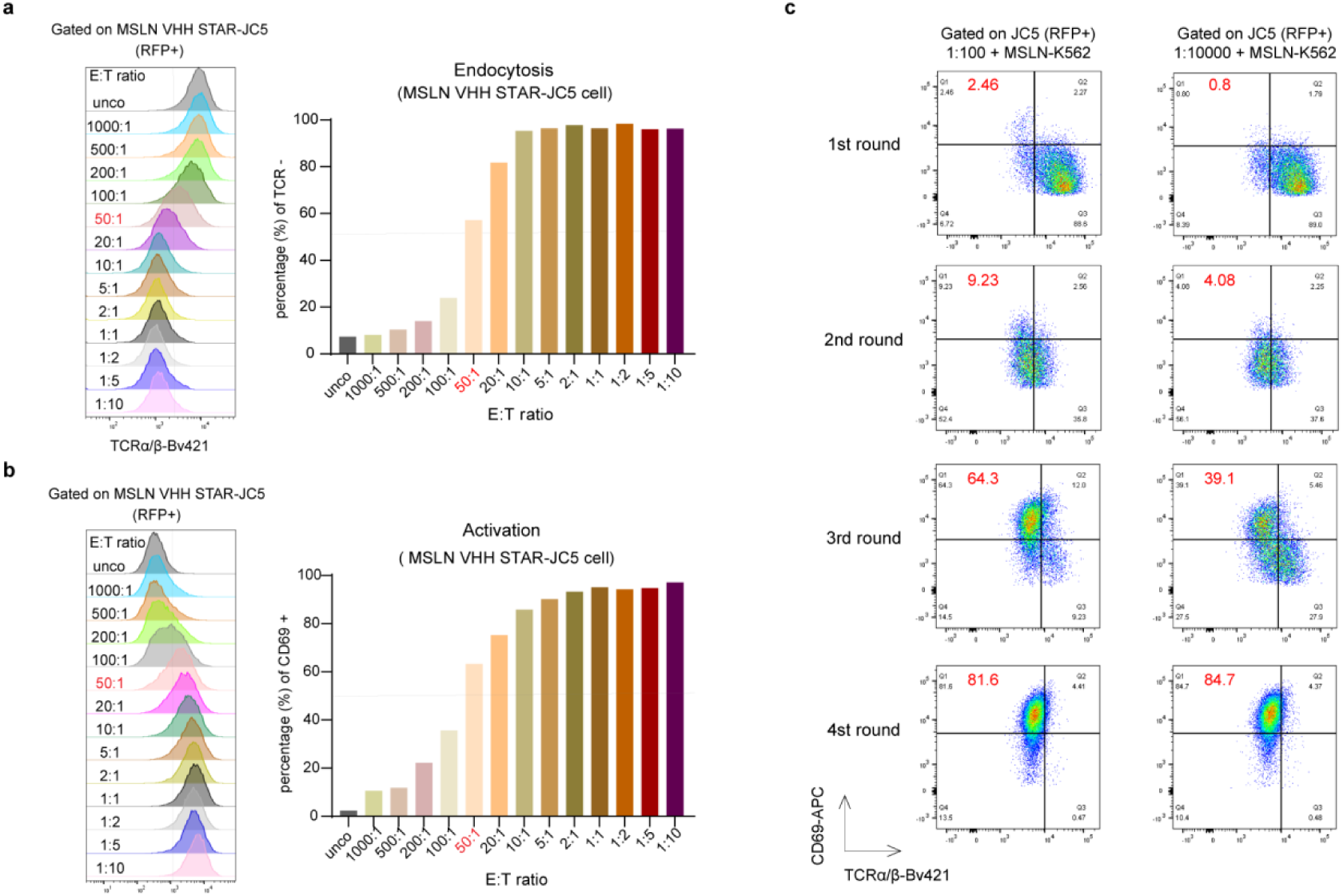
E-A functional index is titratable and sensitive to identify one cognate antibody from 10,000 non-cognate antibodies library. (**a, b**) Flow cytometry analysis of STAR endocytosis and CD69 activation in MSLN VHH STAR-JC5 cells after co-cultured with target MSLN-K562 cells for 24h under different E:T ratios. “Unco” indicates that JC5 cells were not co-cultured with target cells. (**c**) FACS plots of RFP positive JC5 cells from 1:100 or 1:10,000 mixtures after each round of functional screening co-cultured with target MSLN-K562 cells. Data in (**a**) and (**b**) are representative of three independent experiments.

**Supplementary Figure 6.**
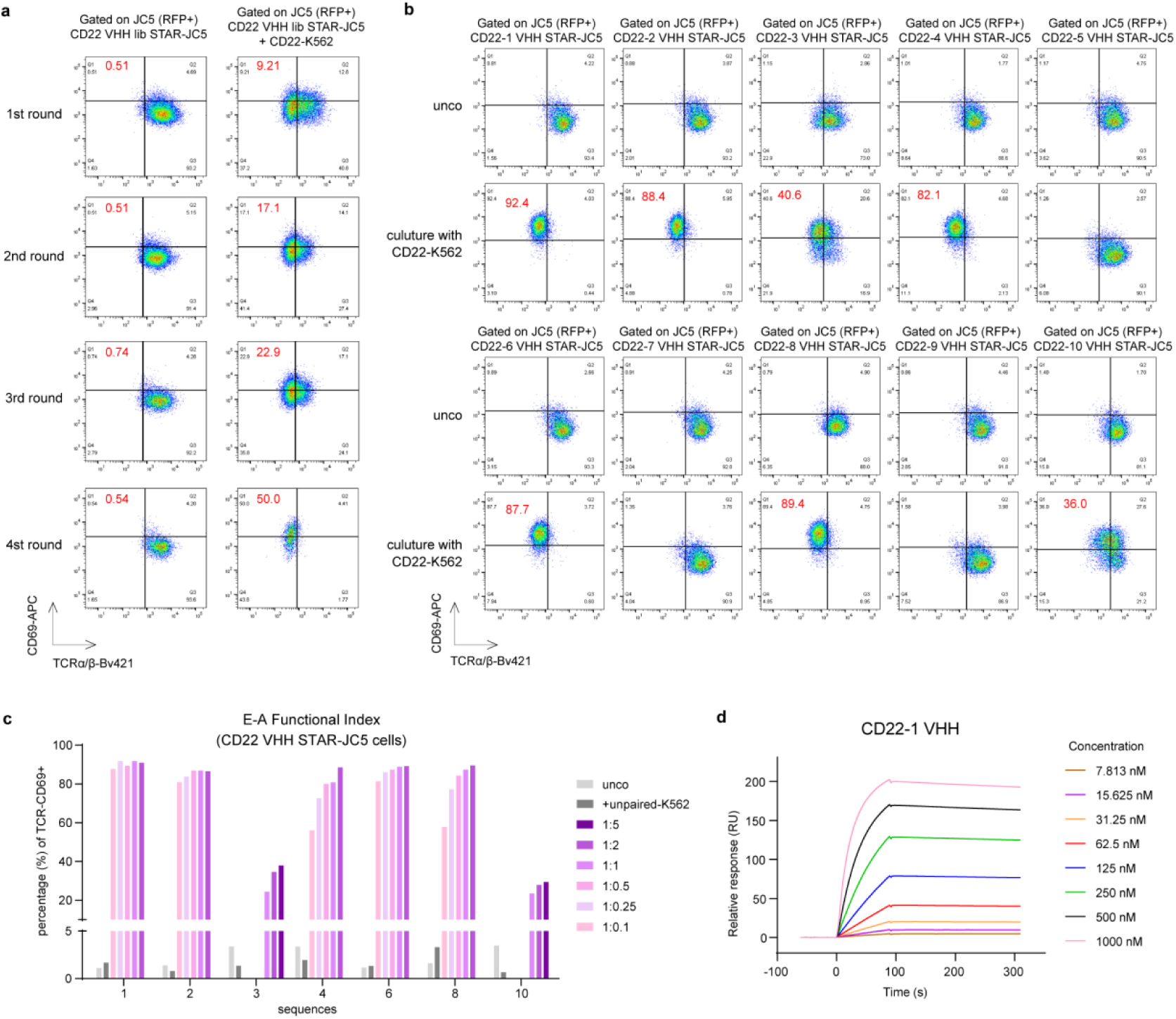
E-A functional screening isolates and identifies membrane antigen CD22-specific antibodies. (**a**) FACS plots of RFP positive JC5 cells from CD22 VHH lib STAR-T cells library after each round of functional screening before and after co-cultured with target CD22-K562 cells (E:T=1:1). (**b**) FACS plots of top ten enriched CD22 VHHs-based STAR-JC5 cells before and after co-cultured with target CD22-K562 cells (E:T=1:1). (**c**) Flow cytometry analysis of seven E-A functional index-positive CD22 VHHs-based STAR-JC5 cells before and after co-cultured with target CD22-K562 cells under different E:T ratios or unpaied-K562 cells as negative control (The results of CD22-3 and CD22-10 groups at E:T ratio of 1:0.5, 1:0.25 and 1:0.1 are not shown). (**d**) CD22-1 VHH (human IgG1-Fc dimer) binding to human CD22 protein was measured with multi cycle kinetics by using SPR. CD22-1 VHH was loaded at increasing concentration and the sensorgram show raw data (coloured by concentration). The co-culture time used in the experiments is 24h. Data are representative of three independent experiments.

**Supplementary Figure 7.**
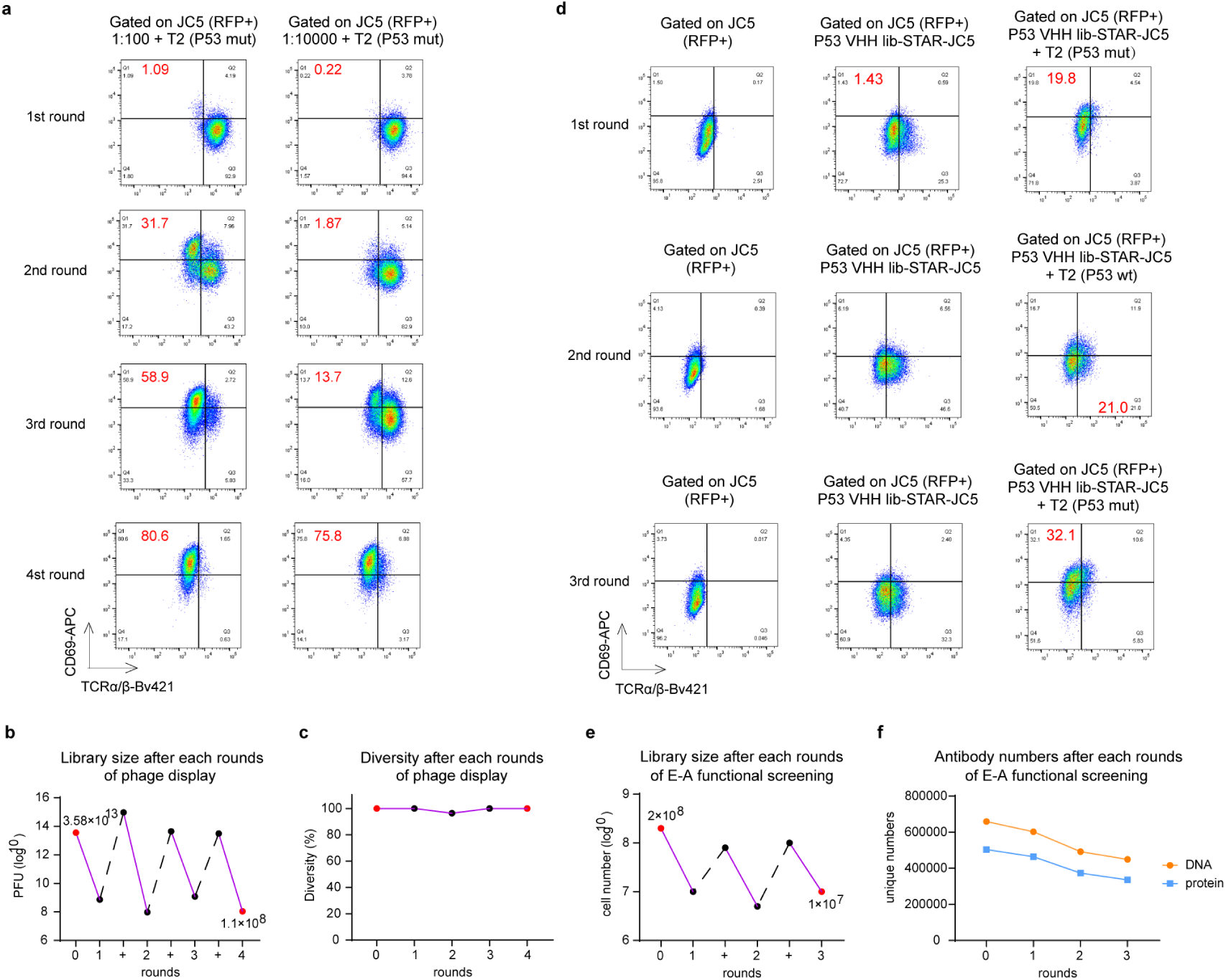
E-A functional screening isolates and identifies neoantigen-specific antibodies. (**a**) FACS plots of RFP positive JC5 cells from 1:100 or 1:10,000 mixtures after each round of functional screening co-culturing with T2 cells loaded P53 mut peptides. Peptides concentration used in each round is 10 µM. (**b, c**) The phage library size (plaque forming units, PFU) and diversity (%) before (0 round) and after four rounds of phage pre-enrichment was measured by randomly selecting 30 clones from the phage library and performing DNA sequencing and sequence comparison analysis.The plus signs on the horizontal axis and black dotted lines in the graph represent amplication stages after each round of bio-panning. (**d**) FACS plots of gated P53 VHH lib STAR-JC5 cells after each round of functional screening. Target cells used in the first and third rounds screening are T2 cells loaded P53 mut peptides, and the second round used P53 wt peptides. Peptides concentration used in each round is 1 µM. (**e**) The library size (cell number) of P53 VHH lib-STAR-JC5 cells before (0 round) and after three rounds of E-A functional screening was measured by FACS sorting of TCR-positive population. The plus signs on the horizontal axis and black dotted lines in the graph represent amplication stages after each round of screening. (**f**) Unique DNA and protein numbers of P53 VHH lib before (0 round) and after each round of E-A functional screening were detected by NGS.

**Supplementary Figure 8.**
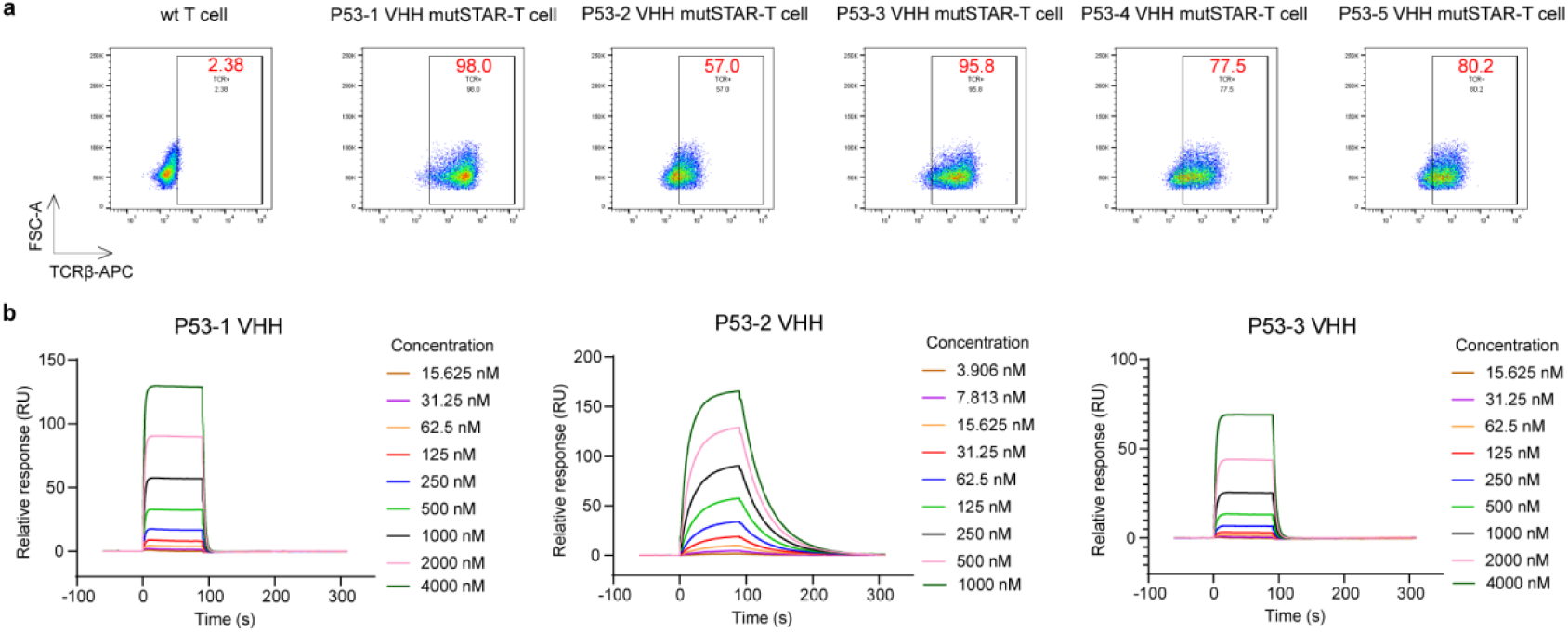
Surface display and antigen-binding of identified P53 VHHs. (**a**) Surface display efficiency of different P53 VHH mutSTAR constructs on primary T cells was assessed by flow cytometry. FSC were used to define the single cell population, the expression of mutSTAR was defined by staining with anti-mouse TCRβ-APC antibody and the positive population was selected. (**b**) Top three P53 VHHs (human IgG1-Fc dimer) binding to P53 R175H-HLA-A*02:01 monomer was measured with multi cycle kinetics by using SPR. VHHs were loaded at increasing concentration respectively and the sensorgram show raw data (coloured by concentration). Data are representative of three independent experiments.

**Supplementary Table 1.** Sequence of VHHs identified from functional screening platform in this study.

| ID | VHH amino acid sequence (CDR3 is highlighted) |
| --- | --- |
| CD22-1 | Upon Correspondance |
| CD22-2 | Upon Correspondance |
| CD22-3 | Upon Correspondance |
| CD22-4 | Upon Correspondance |
| CD22-5 | Upon Correspondance |
| CD22-6 | Upon Correspondance |
| CD22-7 | Upon Correspondance |
| CD22-8 | Upon Correspondance |
| CD22-9 | Upon Correspondance |
| CD22-10 | Upon Correspondance |
| P53-1 | Upon Correspondance |
| P53-2 | Upon Correspondance |
| P53-3 | Upon Correspondance |
| P53-4 | Upon Correspondance |
| P53-5 | Upon Correspondance |

## Notes

### Competing Interest Statement

The authors have declared no competing interest.

### Summary of Updates

We have changed the order of author list by moving the Corresponding Author into last position. Sorry for the mistake.

## References

1. Alpaugh, M. & Cicchetti, F. A brief history of antibody-based therapy. Neurobiol Dis 130, 104504 (2019).

2. Crescioli, S. et al. Antibodies to watch in 2024. MAbs 16, 2297450 (2024).

3. Goydel, R.S. & Rader, C. Antibody-based cancer therapy. Oncogene 40, 3655–3664 (2021).

4. Zinn, S. et al. Advances in antibody-based therapy in oncology. Nat Cancer 4, 165–180 (2023).

5. Liu, Y. et al. Chimeric STAR receptors using TCR machinery mediate robust responses against solid tumors. Sci Transl Med 13 (2021).

6. Wang, J. et al. A novel adoptive synthetic TCR and antigen receptor (STAR) T-Cell therapy for B-Cell acute lymphoblastic leukemia. Am J Hematol 97, 992–1004 (2022).

7. Burton, J. et al. Inefficient exploitation of accessory receptors reduces the sensitivity of chimeric antigen receptors. Proc Natl Acad Sci U S A 120, e2216352120 (2023).

8. Kretzschmar, T. & von Ruden, T. Antibody discovery: phage display. Curr Opin Biotechnol 13, 598–602 (2002).

9. Kanamori, T., Fujino, Y. & Ueda, T. PURE ribosome display and its application in antibody technology. Biochim Biophys Acta 1844, 1925–1932 (2014).

10. Miller, K.D., Pefaur, N.B. & Baird, C.L. Construction and screening of antigen targeted immune yeast surface display antibody libraries. Curr Protoc Cytom Chapter 4, Unit4 7 (2008).

11. Jakobovits, A., Amado, R.G., Yang, X., Roskos, L. & Schwab, G. From XenoMouse technology to panitumumab, the first fully human antibody product from transgenic mice. Nat Biotechnol 25, 1134–1143 (2007).

12. Saeed, A.F., Wang, R., Ling, S. & Wang, S. Antibody Engineering for Pursuing a Healthier Future. Front Microbiol 8, 495 (2017).

13. Fitzgerald, V. & Leonard, P. Single cell screening approaches for antibody discovery. Methods 116, 34–42 (2017).

14. Sun, H., Hu, N. & Wang, J. Application of microfluidic technology in antibody screening. Biotechnol J 17, e2100623 (2022).

15. Pedrioli, A. & Oxenius, A. Single B cell technologies for monoclonal antibody discovery. Trends Immunol 42, 1143–1158 (2021).

16. Shembekar, N., Hu, H., Eustace, D. & Merten, C.A. Single-Cell Droplet Microfluidic Screening for Antibodies Specifically Binding to Target Cells. Cell Rep 22, 2206–2215 (2018).

17. Shapiro, M.B. et al. Alpaca single B cell interrogation and heavy-chain-only antibody discovery on an optofluidic platform. Antib Ther 6, 211–223 (2023).

18. Bennett, N.R. et al. Atomically accurate de novo design of single-domain antibodies. bioRxiv (2024).

19. Bloemberg, D. et al. A High-Throughput Method for Characterizing Novel Chimeric Antigen Receptors in Jurkat Cells. Mol Ther Methods Clin Dev 16, 238–254 (2020).

20. Ma, P. et al. Avidity-Based Selection of Tissue-Specific CAR-T Cells from a Combinatorial Cellular Library of CARs. Adv Sci (Weinh) 8, 2003091 (2021).

21. Charpentier, J.C. & King, P.D. Mechanisms and functions of endocytosis in T cells. Cell Commun Signal 19, 92 (2021).

22. Evnouchidou, I., Caillens, V., Koumantou, D. & Saveanu, L. The role of endocytic trafficking in antigen T cell receptor activation. Biomed J 45, 310–320 (2022).

23. Cibrian, D. & Sanchez-Madrid, F. CD69: from activation marker to metabolic gatekeeper. Eur J Immunol 47, 946–953 (2017).

24. Peri, A. et al. The landscape of T cell antigens for cancer immunotherapy. Nat Cancer 4, 937–954 (2023).

25. Hsiue, E.H. et al. Targeting a neoantigen derived from a common TP53 mutation. Science 371 (2021).

26. Chai, D. et al. DNA-delivered monoclonal antibodies targeting the p53 R175H mutant epitope inhibit tumor development in mice. Genes Dis 11, 100994 (2024).

27. Liu, M., Li, L., Jin, D. & Liu, Y. Nanobody-A versatile tool for cancer diagnosis and therapeutics. Wiley Interdiscip Rev Nanomed Nanobiotechnol 13, e1697 (2021).

28. Bao, C. et al. The Application of Nanobody in CAR-T Therapy. Biomolecules 11 (2021).

29. Carter, P.J. & Rajpal, A. Designing antibodies as therapeutics. Cell 185, 2789–2805 (2022).

30. t Hoen, P.A. et al. Phage display screening without repetitious selection rounds. Anal Biochem 421, 622–631 (2012).

31. Alonso-Camino, V. et al. CARbodies: Human Antibodies Against Cell Surface Tumor Antigens Selected From Repertoires Displayed on T Cell Chimeric Antigen Receptors. Mol Ther Nucleic Acids 2, e93 (2013).

32. Xie, N. et al. Neoantigens: promising targets for cancer therapy. Signal Transduct Target Ther 8, 9 (2023).

33. Schumacher, T.N. & Schreiber, R.D. Neoantigens in cancer immunotherapy. Science 348, 69–74 (2015).

34. Zhang, Z. et al. Neoantigen: A New Breakthrough in Tumor Immunotherapy. Front Immunol 12, 672356 (2021).

35. He, Q. et al. TCR-like antibodies in cancer immunotherapy. J Hematol Oncol 12, 99 (2019).

36. Xie, G. et al. CAR-T cells targeting a nucleophosmin neoepitope exhibit potent specific activity in mouse models of acute myeloid leukaemia. Nat Biomed Eng 5, 399–413 (2021).

37. Douglass, J., et al. Bispecific antibodies targeting mutant RAS neoantigens. Sci Immunol 6 (2021).

38. Huang, D. et al. TCR-mimicking STAR conveys superior sensitivity over CAR in targeting tumors with low-density neoantigens. Cell Rep 43, 114949 (2024).

39. Nicholson, I.C. et al. Construction and characterisation of a functional CD19 specific single chain Fv fragment for immunotherapy of B lineage leukaemia and lymphoma. Mol Immunol 34, 1157–1165 (1997).

40. Nakano, K. et al. Generation of a humanized anti-glypican 3 antibody by CDR grafting and stability optimization. Anticancer Drugs 21, 907–916 (2010).

41. Van Wauwe, J.P., De Mey, J.R. & Goossens, J.G. OKT3: a monoclonal anti-human T lymphocyte antibody with potent mitogenic properties. J Immunol 124, 2708–2713 (1980).

42. Wei, H. et al. Structural basis of a novel heterodimeric Fc for bispecific antibody production. Oncotarget 8, 51037–51049 (2017).

